# Traditional functional groups capture limited variation in the trait space of macroalgae

**DOI:** 10.1101/803965

**Authors:** Alizée R. L. Mauffrey, Laura Cappelatti, John N. Griffin

## Abstract

1. Macroalgal (seaweed) beds and forests fuel coastal ecosystems and are rapidly reorganising under global change, but quantifying their functional structure still relies on binning species into coarse groups on the assumption that they adequately capture relevant underlying traits.
2. To interrogate this “group gambit”, we first measured 12 traits relating to competitive dominance and resource economics across 95 macroalgal species collected from UK rocky shores. We then assessed trait variation explained by traditional grouping approaches consisting of (i) two highly-cited schemes based on gross morphology and anatomy and (ii) two commonly-used categorisations of vertical space use. To identify the limitations of traditional grouping approaches and to reveal potential alternatives, we also assessed the ability of (iii) emergent groups created from *post hoc* clustering of our dataset to account for macroalgal trait variation.
3. (i) Traditional groups explained about a third of multivariate trait expression with considerable group overlap. (ii) Classifications of vertical space use accounted for even less multivariate trait expression. Notwithstanding considerable overlap, the canopy vs. turf scheme explained significant differences in most individual traits, with turf species tending to display attributes of opportunistic forms. (iii) Emergent groups were substantially more parsimonious than all existing grouping approaches.
4. Synthesis: Our analysis using a comprehensive dataset of directly measured functional traits failed to strongly support the group gambit in macroalgae. While existing grouping approaches may allow first order approximations, they risk considerable loss of information at the trait and, potentially, ecosystem levels. We call for further development of a trait-based approach to macroalgal functional ecology to capture unfolding community and ecosystem changes with greater accuracy and generality.

## 1. INTRODUCTION

Macroalgae (seaweed) form extensive and productive beds in the coastal environment worldwide (Hurd, Harrison, Bischof, & Lobban, 2014). These phylogenetically and functionally diverse foundation species fuel coastal ecosystems and provide a range of ecosystem services including food and natural products (Griffiths, Harrison, Smit, & Maharajh, 2016; McLachlan, 1985), habitat for commercially important fish (Miranda, Lopez-Alonso, & Garcia-Vaquero, 2017), and blue carbon sequestration (Chung, Beardall, Mehta, Sahoo, & Stojkovic, 2011). Yet under accelerating anthropogenic forcing, macroalgal beds are experiencing major changes in community composition as, for instance, kelp retreat towards the poles while warmer-water and non-native species expand (Harley et al., 2012; Pessarrodona, Foggo, & Smale, 2019). Such community restructuring – and its ecosystem ramifications – is mediated by species’ functional traits (Lavorel & Garnier, 2002; Suding et al., 2008; Trugman et al., 2019).

Traits hold the promise of more predictive ecology that goes beyond case studies of species or taxonomic groups towards a more general understanding (McGill, Enquist, Weiher, & Westoby, 2006). Functional traits are morphological, physiological or phenological characteristics of individuals that influence their response to the environment and/or effect on ecosystem properties and/or services (Díaz et al., 2013). Traits reflect adaptive strategies and underlying physiological tradeoffs. In vascular plants, for example, trait variation largely reflects competitive dominance (plant size) and the leaf economics spectrum, which emerges from physiological tradeoffs between structural integrity and growth potential (Díaz et al., 2016). As an alternative to direct use of trait values, so-called ‘emergent’ groups have been built through *post hoc* clustering of trait data on the premise that they represent trait variability more closely than traditional grouping approaches (Lavorel, McIntyre, Landsberg, & Forbes, 1997). In turn, functional traits or emergent groups explain species’ contributions to ecosystem functions and services. For instance, plant specific leaf area and leaf nitrogen content have been linked to primary productivity and decomposition (Reich, 2014; Shipley, Lechowicz, Wright, & Reich, 2006). Terrestrial vascular plants have an inordinate influence on the development of functional ecology of autotrophic organisms, but efforts are underway to include primary producers from lichens and bryophytes to aquatic macrophytes such as seagrasses and freshwater plants (de los Santos et al., 2016; Elger & Willby, 2003; Roos et al., in press).

However, ecologists studying macroalgal beds and forests rarely directly measure functional traits and instead assign species to long-established form-centric groups. Littler and Littler (1980) first proposed the ‘functional-form model’, categorising species into morpho-functional groups based on gross morphology and anatomical features. Over a decade later, Steneck and Dethier (1994) proposed a less subjective classification based on branching pattern, anatomy and degree of cortication. Macroalgal ecologists have also employed broad categorisations of vertical space use, such as the binary canopy vs. turf scheme, to infer ecosystem consequences of changing assemblage structure, particularly in light of the global rise of turfs over canopy-forming macroalgae (Feehan, Grace, & Narvaez, 2019; Filbee-Dexter & Wernberg, 2018). Traditional grouping methods remain highly influential: for instance, Steneck and Dethier’s paper has been cited over 1000 times, with 60 citations in 2018 alone (Google Scholar, accessed October 2019).

Macroalgal groups are much less labour-intensive than direct trait measurements but carry implicit assumptions. Grouping approaches overlook within-group variability (Petchey & Gaston, 2006), can lead to subjective species allocation (Chapin, Bret-Harte, Hobbie, & Zhong, 1996; Díaz & Cabido, 2001; Phillips, Kendrick, & Lavery, 1997), and disregard relative influences of specific traits and intraspecific variability (Hanisak, Littler, & Littler, 1988; Padilla & Allen, 2000; Violle et al., 2012). Perhaps most importantly from the ecosystem perspective, it is assumed that macroalgal groups capture relevant suites of functional traits and, in turn, ecological processes. For instance, ‘sheet species’ are expected to possess a high relative surface area, allowing fast nutrient acquisition and high productivity (Littler & Littler, 1980; Steneck & Dethier, 1994) while members of the canopy typically display high structural integrity and complexity, thereby sustaining rich epibiota (Filbee-Dexter & Wernberg, 2018; Teagle, Hawkins, Moore, & Smale, 2017). The assumption that traditional macroalgal functional classifications capture underlying interspecific trait variation remains untested. We call this ‘the group gambit’ (following the phylogenetic hypothesis in Mazel et al. 2018).

Here, we provide the first comprehensive test of the group gambit in macroalgae. Using direct measurements of traits related to two major aspects of ecological variation, competitive dominance and resource economics, we test the assumption that grouping approaches explain a large proportion of interspecific trait variation across four commonly-used traditional schemes: the form-centric (i.e., morphology and anatomy-based) grouping approaches of Littler and Littler (1) and Steneck and Dethier (2), as well as two classifications of macroalgal vertical space use, or stature: the binary canopy vs. turf (3) and a three-level scheme adapted from Arenas, Sánchez, Hawkins, and Jenkins (2006; 4). To further probe the validity of traditional groups, we also assess how *post hoc* clustering of our dataset into emergent groups leads to reclassification of species and accounts for macroalgal functional variation. A strong correspondence between groups and the traditional underlying functional traits would support their continued application and bolster ecological interpretation. However, substantial mismatch between groups and traits – as well as reclassification of species into emergent groups – would indicate a loss of information and underline the potential gains offered by direct trait measurements.

## 2. MATERIALS AND METHODS

### 2.1. Sampling

We measured eleven continuous and one categorical functional traits (Table S1) at the individual level across 95 erect intertidal macroalgal species, which spanned a great variety of form and function and hence, traditional functional groups (Table S2). Samples were collected samples from twelve rocky shores in the UK ranging from very sheltered to very exposed (Table S3). Sampling took place from May to September 2013 and 2015-2018 (Table S4). We collected an average of 6 replicates per species, ranging from 1 to 45 (mode and median = 6, S.D. = 7.55; Table S4). Such a large difference in replication was due to the rarity of some species and to our efforts in sampling abundant species across several sites to better capture natural variability. Replicates were sampled more than 2 m apart. Whenever possible, a replicate was made up of one lone-standing individual. However, when individuals were too small for the trait measurements, a sufficient quantity was collected for each replicate by pooling several individuals or tufts (Table S4). Whenever distinguishing between individuals was not possible (e.g., for turfs), the samples were collected by isolating tufts (Table S4). Samples were kept in seawater in a cooler until brought back to the laboratory. They were then either screened fresh or frozen at −18°C until processed.

### 2.2. Trait screening

The functional traits measured are hypothesised to capture two fundamental aspects of primary producer variability, (1) the economics spectrum and (2) competitive dominance. We consider multiple indicators (or ‘functional markers’ *sensu* Garnier et al., 2004) to provide a more integrated estimation of ecological strategy and function. Here, we briefly summarise the ecological significance of the traits with regard to photosynthesis, structural integrity, space use and complexity (see Tables S1 and S5 for additional information). The suite of economics-related traits indicates the relative investment in resource acquisition versus resistance to (a)biotic stress and therefore resource conservation, tying in with the *r*- (‘fast return’) to *K*- (‘slow-return’) selection continuum (Pianka, 1970). Slow-return primary producers tend to display long lifespans, low maximum photosynthesis and productivity, reduced palatability, and slow decomposition (Littler & Littler, 1980; Smart et al., 2017; Wright et al., 2004). Traits *a-g* relate to photosynthesis and/or structural integrity, and hence, position on the economics spectrum: Thallus Dry Matter Content (TDMC; a) is the ratio between dry and wet mass and represents the proportion of structural compounds and water-filled – and therefore mainly photosynthetically active – tissues (Elger & Willby, 2003; Littler & Littler, 1981; Schonbeck & Norton, 1979). Frond thickness (b) also increases with the amount of structural tissue, providing resistance to physical stress and herbivore grazing (Cappelatti, Mauffrey, & Griffin, 2019; Littler & Littler, 1980; Littler, Taylor, & Littler, 1983); Carbon content (c) and its ratio (d) with Nitrogen content (e) more directly quantify recalcitrant structural compounds relative to N-rich photosynthetically-active tissues (Cornelissen et al., 2003; Weykam, Gómez, Wiencke, Iken, & Klöser, 1996). Specific Thallus Area (STA; f), analogously to Specific Leaf Area (Wilson, Thompson, & Hodgson, 1999), captures light- and nutrient-absorbing surfaces and increases with the extent of low density, water-filled, photosynthetically-active tissues relative to recalcitrant, structural compounds (Littler & Littler, 1980). Finally, because macroalgae absorb nutrients through the blades, SA:V (g) is associated with nutrient acquisition (Littler & Littler, 1980).

Traits *h-l* are hypothesised to relate to space use and complexity, and hence, competitive dominance. Plant height is a major determinant of competitive dominance (Díaz et al., 2016). Its macroalgal analogue, maximum length (h), and by extension aspect ratio (i), relates to the ability of macroalgae to outcompete surrounding individuals by taking the position of canopy and emerges from a tradeoff between light capture and structural integrity (Carpenter, 1990; Littler & Littler, 1980). The presence of pneumatocysts (j) is another important predictor of macroalgal competitive dominance through canopy occupancy (Dromgoole, 1981). Branching order (k) and SA:P (l) relate to three-dimensional complexity and resource acquisition, and hence, both competitive dominance and economics (Steneck & Dethier, 1994; Veiga, Rubal, & Sousa-Pinto, 2014). High complexity allows individuals to maximise light exposure (Stewart & Carpenter, 2003), provides greater nutrient and gas exchange, delays desiccation at low tide (Hay, 1981; Padilla, 1984; Taylor & Hay, 1984) and reduces the impact of herbivory (Padilla, 1984), but increases drag (Starko, Claman, & Martone, 2015). We measured both traits at the whole individual level to capture the complexity of the whole thallus, since all thallus parts, from holdfast to fronds, are important habitats for epibiota and nekton (Teagle et al., 2017).

Large or structurally complex individuals were subsampled, ensuring that the proportions of all thallus parts were respected (Table S4). We measured the surface area and perimeter of partly microscopic species on subsamples under the microscope and proportionally scaled them up to the whole sample (Table S4). The samples were cleaned of epibiota in seawater and rinsed in deionised water for elemental screening. To obtain TDMC, we recorded sample wet and oven-dried mass (g; Ohaus Scout Pro SP402, SPU602 and Pioneer Analytical PA114, Parsippany, NJ, USA). Thickness (mm) was averaged from ten measurements taken haphazardly along the fronds (Digital Micrometers Ltd, DTG03 0.005, DML3032 0.001 mm, Sheffield, UK), avoiding when applicable the midrib. Individuals were displayed on a lightbox (MiniSun A1, Manchester, UK) and photographed (Pentax K3 digital camera, SMC DA L 18-55 mm, Tokyo, Japan), scanned (Epson Perfection V600, V39, Suwa, Japan), or photographed with an imaging microscope (Leica S8AP0, Wetzlar, Germany, affixed with GT Vision GXCAM-H3, Sudbury, UK). We measured frond (when differentiated) or whole-individual surface area (mm^2^) and whole-individual perimeter (mm) using the software ImageJ (Schneider, Rasband, & Eliceiri, 2012), and calculated SA:V (mm^2^ mL^−1^), STA (mm^2^ g^−1^), and SA:P. Volume (mL) was measured by water displacement. To obtain C and N content and C:N, powderised samples were run through an elemental analyser.

### 2.3. Categorisation of species into functional groups

We allocated species to the groups defined by Littler and Littler as well as Steneck and Dethier based on a review of the literature (Table S2). The species we screened belonged to five traditional groups: ‘filamentous’, ‘sheet’, ‘coarsely branched’, ‘thick leathery’ and ‘articulate calcareous’ for Littler and Littler’s functional-form model, and five groups defined by Steneck and Dethier (1994), ‘filamentous (S)’, ‘foliose’, ‘corticated’, ‘leathery’ and ‘articulated calcareous’, in increasing order of cortication. Although both schemes contain groups with similar or identical names, they were originally defined using different approaches and are not assumed *a priori* to be analogous. We also tested Steneck and Dethier’s detailed classification, which includes two subgroups (Supplementary information, section 1). We used two common classifications of vertical space use: the binary canopy vs. turf scheme and a three-level canopy/subcanopy/turf classification adapted from Arenas et al. (2006). Turfs were considered macroalgae with little to no three-dimensional structure compared with kelp and other canopy-forming macroalgae that form a dense layer of fine filaments, branches, or plumes on the substratum (Filbee-Dexter & Wernberg, 2018). This broad definition of turf macroalgae allowed classification of all species within our study. Location along the canopy is somewhat community-dependent, so we categorised species into the three-level scheme based on what we judged was the most common scenario on the rocky shores screened.

### 2.4. Data analysis

We performed all analyses on R 3.5.3 (R Core Team, 2019) and plotted graphical results using ggplot2 (Wickham, 2009) and ggpubr (Kassambara, 2019). Prior to running analyses on species-level traits, we examined individual-level trait variability through boxplots and Kruskal-Wallis one-way analyses of variance (Fig. S1). Species trait averages were transformed to bring their distribution as close to normality as possible and to reduce differences in scale across traits (Table S5). To examine the distribution of individual traits among traditional groups, smoothed density curves for each of the eleven continuous functional traits studied were drawn. To assess whether groups differed from each other, we ran pairwise Wilcoxon signed-rank tests on every group pair.

We imputed the four percent of average trait values that were missing from the dataset (function “mice” in eponymous R package; (van Buuren & Groothuis-Oudshoorn, 2011) to then reduce the dimensionality of the data using a Principal Coordinate Analysis (PCoA; function “cmdscale”). A sensitivity analysis revealed that the PCoA scores and those of a Principal Component Analysis (PCA) run on the same dataset (minus the binary trait ‘pneumatocysts’), i.e., the general position of species in trait space, were affected to some extent by the dimensionality reduction method (Fig. S2). We favoured a PCoA because it allowed us to include pneumatocyst presence and because it accounted for underlying trait correlations without assuming linearity, which many of the trait pairs violated (Table S6). The PCoA was run on a weighted Gower matrix (function “daisy” in “cluster”; Maechler, Rousseeuw, Struyf, Hubert, & Hornik, 2019), with equal weighting to traits associated with photosynthesis, structural integrity, space use, and complexity (Table S5). To assess the strength of association between the principal coordinates and each trait, we ran linear regressions between the scores of the first three PCoA axes and the twelve traits studied.

Two clustering methods were used to create emergent groups from the weighted Gower matrix: agglomerative Hierarchical Cluster Analysis (HCA) and *k*-medoids clustering (*k*-medoids). Agglomerative HCA performs bottom-up clustering of species by iteratively combining groups with similar traits (Lukasová, 1979; using Ward’s minimum variance or “ward.D2” in “hclust”; (Murtagh & Legendre, 2014). *K*-medoids is a top-down clustering approach whereby species are assigned to a chosen number of groups based on multivariate distance from group medoids, making it rather robust to noise and outliers (Reynolds, Richards, de la Iglesia, & Rayward-Smith, 2006; function “pam” of package “cluster”, Maechler et al., 2019). We ran each clustering method using five clusters to match the number of clusters used by the functional-form model and Steneck and Dethier’s classification. To assess the extent of trait variance explained by traditional and emergent groups, we ran a series of PERMANOVAs on the weighted Gower matrix using the grouping approaches studied as a grouping variable. The PERMANOVAs were run with 999 permutations (“adonis” function in “vegan”; Oksanen et al., 2019). To compare support for PERMANOVA models with different numbers of groups, we calculated pseudo-AIC values using residual sum of squares.

## 3. RESULTS

### 3.1. Distributions of individual traits

All traits displayed greater inter- than intra-specific variability (Fig. S1). All results reported hereafter are based on species trait averages.

For both traditional grouping approaches, trait distributions were generally not unimodal and were right-skewed, suggesting that while many species shared similar trait values within a group, several others were functionally distinct and groups poorly captured natural variation (Fig. 1; see Fig. S3 for Steneck and Dethier’s classification). Under both schemes, group overlap was extensive across traits; from a possible 55 cases (11 traits × 5 groups), there were only 7 instances where a group was different from all others (i.e., functionally unique; Fig. 1, S3). None of the form-related groups were functionally unique for thickness, aspect ratio, branching order, SA:P, or C:N. However, under both schemes, there were differences between roughly half of all possible group pairs, reflecting the position of groups along the continuum of trait variability (Figs 1, S1, S4-S5). Overall, ‘(thick) leathery’ and ‘articulate(d) calcareous’ were the two most distinguished functional groups (Figs 1, S3).

**Figure 1.**
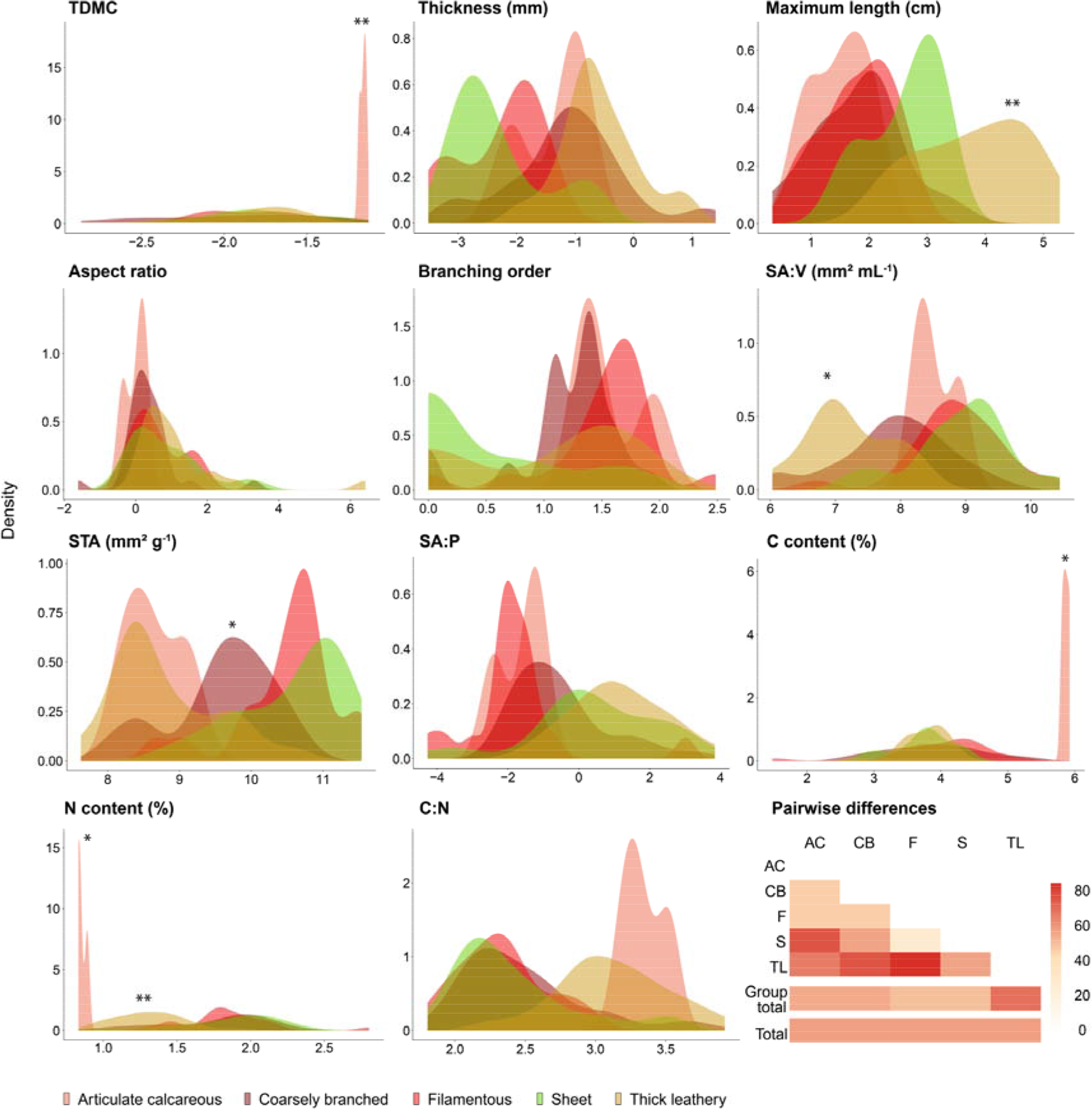
Distribution of species-level traits among Littler and Littler’s groups. Values of the 11 continuous functional traits are transformed species-level means across the 95 macroalgal species screened (Table S5). Groups that are significantly different from all others are marked by asterisks (pairwise Wilcoxon rank sum test; ‘*’: *P* < 0.05, ‘**’: *P* < 0.01; lowest common *P* is shown). Heatmap (bottom right) shows the proportion of significant differences among pairwise comparisons for each group (group total) and overall (total; ‘AC’ is for articulated calcareous, ‘CB’ for coarsely branched, ‘F’ for filamentous, ‘S’ for sheet, and ‘TL’ for thick leathery); see Fig. S4 for pairwise differences among groups for each individual trait.

Both classifications of macroalgal stature explained significant differences in most of the traits’ distributions (Fig 2, see Fig S7 for the three-level classification). Specifically, in the canopy vs. turf scheme, canopy species had greater thickness, maximum length, aspect ratio and C:N, while turf species had greater STA, SA:V, C and N values. In the three-level scheme, all the groups differed for maximum length, thickness, SA:V and STA. Canopy also had lower values than turf for SA:P and N, and greater values for C:N (Figs S6-S8). Notwithstanding these differences, stature-centric groups spanned wide ranges of trait values and in most cases, displayed a high degree of overlap (Fig. 2, S6-8). The prevalence of significant differences between groups, compared to the two form-centric approaches, should be interpreted in light of the higher within-group sample sizes.

**Figure 2.**
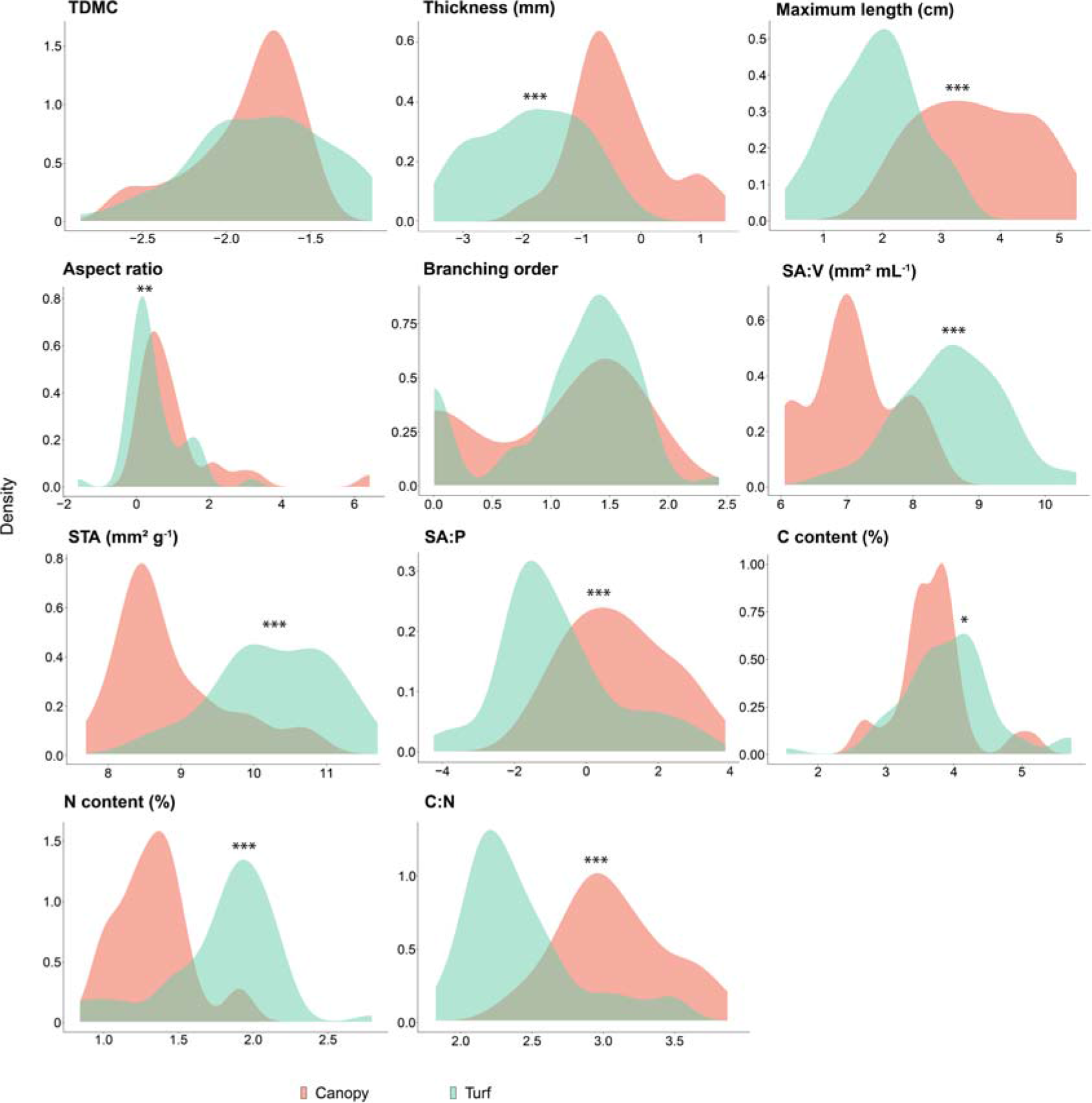
Distribution of species-level traits among a common binary classification of vertical space use. Values of the 11 continuous functional traits are transformed species-level means across the 95 macroalgal species screened (Table S5). Groups that are significantly different from all others are marked by asterisks (pairwise Wilcoxon rank sum test; ‘*’: *P* < 0.05, ‘**’: *P* < 0.01, ‘***’: *P* < 0.001; lowest common *P* is shown.

### 3.2. Distributions in multivariate trait space

Many of the functional traits were entrained along the first PCoA axis. Species positioned further along the first PCoA axis had lower maximum length, C:N ratio and thickness, and higher SA:V, STA and N content (Fig. 3). These trait attributes correspond to a less (light-) competitive but a faster (resource acquisitive) strategy. Meanwhile, a single functional trait (branching order) was clearly most strongly associated with the second PCoA axis. The first two principal coordinates accounted for 49.9% of inertia in the distance matrix, while the third (not shown) accounted for an additional 13%.

**Figure 3.**
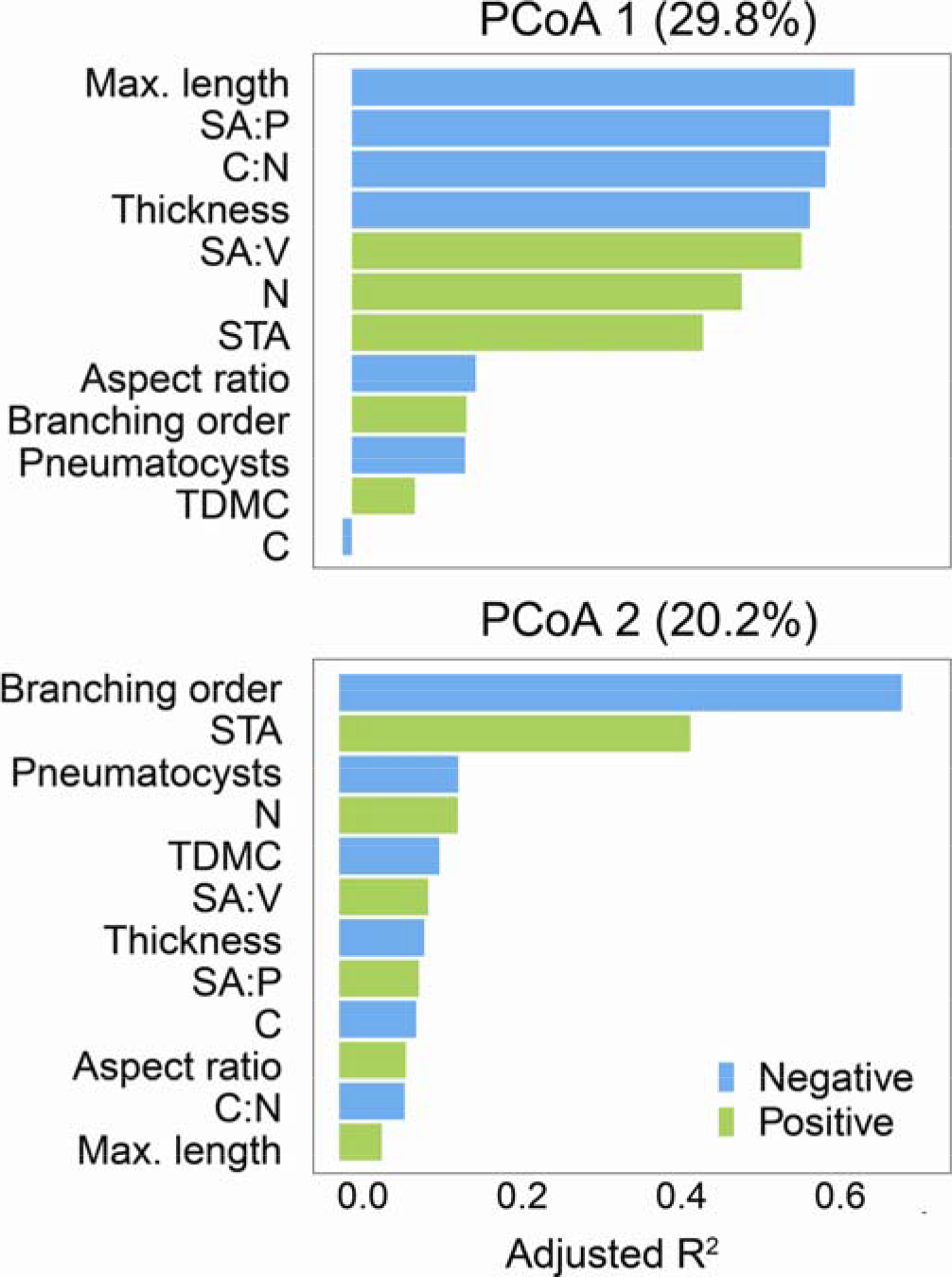
Associations between functional traits and principal coordinate axes. Associations were assessed by a series of linear regressions based on the sample size of 95 species. The strength and sign of trait-axis association are indicated, respectively, by the coefficients of determination (R^2^) and colour of the bars. Proportion of inertia accounted for by each axis is given in parenthesis.

Neither of the grouping methods based on macroalgal form captured a large proportion of dispersion in trait space. Littler and Littler’s and Steneck and Dethier’s groups explained about 37% of multivariate species-level trait expression (Fig. 4, five-cluster PERMANOVA, *R*^2^ = 0.37, *P* < 0.001 and *R*^2^ = 0.37, *P* < 0.001, respectively) while not differing in their level of parsimony (AICs = 63.61 and 63.63). Littler and Littler’s and Steneck and Dethier’s groups were located in roughly the same locations of the trait space (Fig. 4). Species categorised under (thick) leathery displayed a wide array of trait values and were split between those characterised by high SA:P and maximum length on the one hand, and high thickness and C:N on the other (Figs 1 and 4). Calcareous species displayed high C content and branching order, and stood out from the rest of the species. Coarsely branched/corticated species were scattered across most of the trait space. The filamentous groups mainly gathered species with high SA:V and branching order. Finally, the sheet/foliose group principally represented species with low C:N and high STA (Figs 1 and 4).

**Figure 4.**
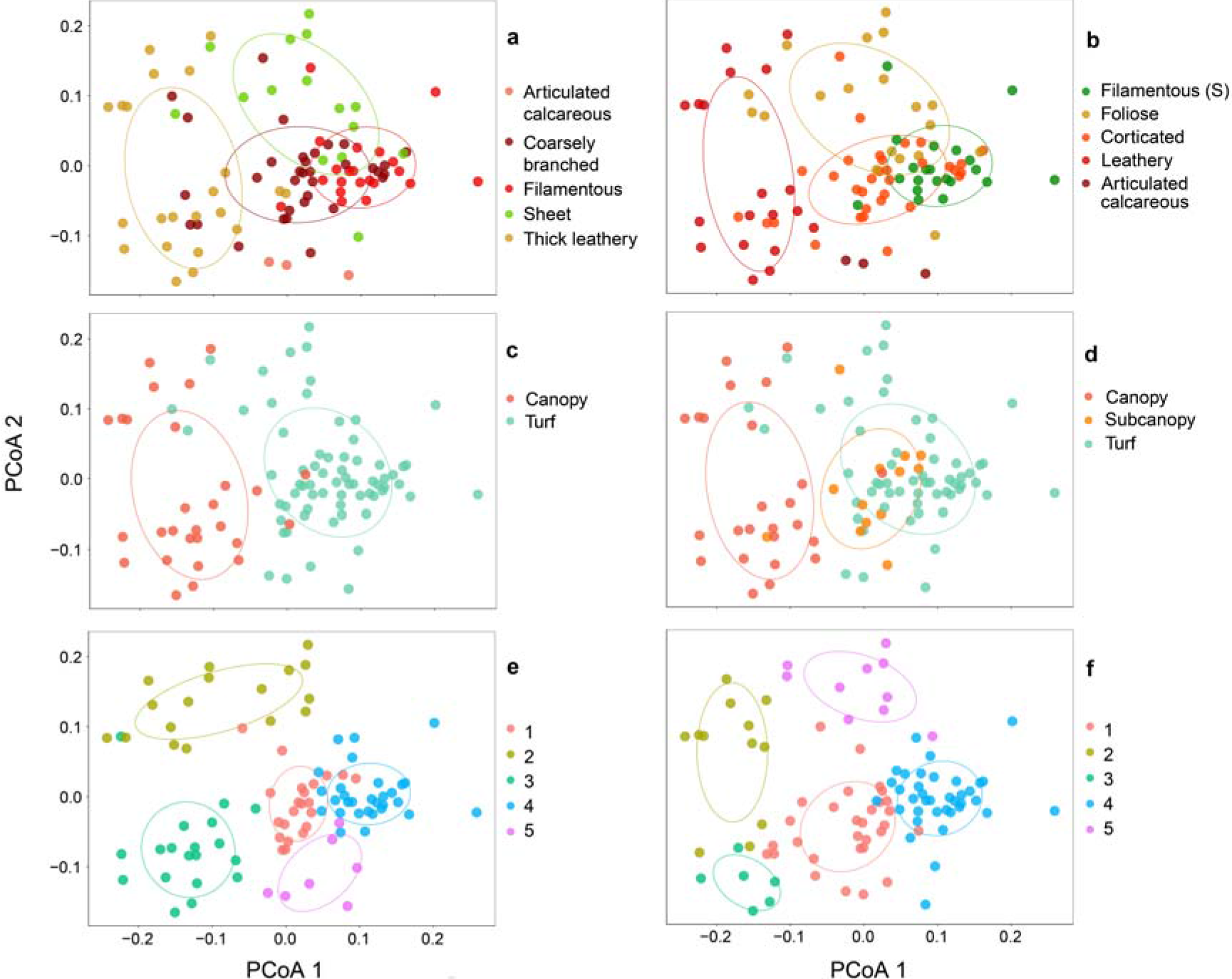
Distribution of functional groups in macroalgal trait space. Group distribution in trait space as yielded by the first two axes of a principal coordinate analysis is given for: (a) Littler and Littler’s ‘functional-form’ model (ordered alphabetically), (b) Steneck and Dethier’s classification (ordered by degree of cortication), (c) canopy vs. turf, (d) canopy/subcanopy/turf, and emergent groups yielded by *post hoc* clustering of the species using (e) agglomerative Hierarchical Clustering Analysis (HCA) and (f) the divisive *k*-medoids method. Confidence ellipses (50%) are represented, assuming a multivariate normal distribution.

Classifications of vertical space use explained little multivariate trait expression and were the least parsimonious of all traditional schemes (versus Littler and Littler’s groups, canopy vs. turf: ΔAIC = 9.07; canopy/subcanopy/turf: ΔAIC = 10.7). The binary canopy vs. turf scheme explained about 26% of multivariate trait expression (two-cluster PERMANOVA, *R*^2^ = 0.26, *P* < 0.001) and – despite containing an extra group – the three-level scheme explained roughly the same amount of trait variation, ca. 27% (three-cluster PERMANOVA, *R*^2^ = 0.26, *P* < 0.001).

Emergent classifications captured more distinct trait information than traditional ones, as seen as more separate clusters in trait space (Fig. 4), and explained greater multivariate trait expression. Regardless of the clustering method, emergent groups explained slightly more than half of the trait variance (55% using HCA, *R*^2^ = 0.55, *P* < 0.001 and 54% using *k*-medoids, *R*^2^ = 0.54, *P* < 0.001, five-cluster PERMANOVA). Both emergent classifications were the most parsimonious of all grouping approaches (versus Littler and Littler’s groups: HCA: ΔAIC = −33.8; *k*-medoids: ΔAIC = −30.85).

### 3.3. Reclassification across grouping approaches

To evaluate how species are classified under different approaches (including emergent groups), the group membership of individual species can be traced across the grouping schemes (Fig. 5). Despite the less subjective basis of Steneck and Dethier’s scheme, largely the same sets of species remained grouped together compared to Littler and Littler’s earlier scheme (Table S7). Correspondences were evident between groups: the ‘filamentous’ group of Littler and Littler’s model corresponded to ‘filamentous (S)’ in Steneck and Dethier’s classification, ‘sheet’ to ‘foliose’, ‘coarsely branched’ to ‘corticated’, ‘thick leathery’ to ‘leathery’, and ‘articulate calcareous’ to ‘articulated calcareous’. All of the mismatches in species allocation between the two schemes were due to Littler and Littler’s groups ‘coarsely branched’ and ‘thick leathery’ being reclassified by Steneck and Dethier’s approach (Fig. 5). The addition of an extra category to the canopy vs. turf scheme, to form canopy/subcanopy/turf, led to the reclassification of larger turf species into the subcanopy group (11% of all species reclassified). These reclassified species were primarily coarsely branched under Littler and Littler’s scheme.

**Figure 5.**
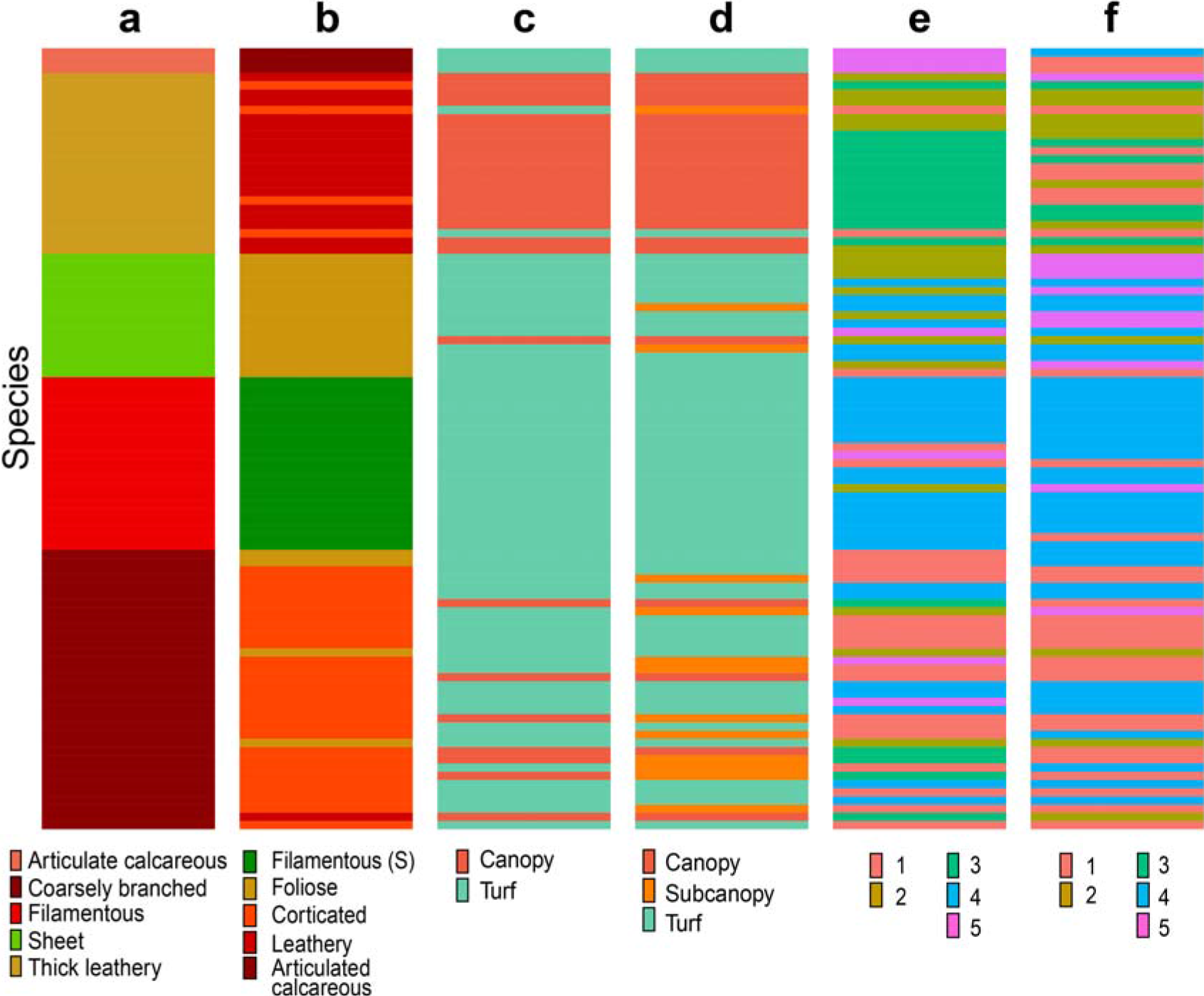
Species re-allocation across grouping approaches. Species composition is given for (a) Littler and Littler’s ‘functional-form’ model (ordered alphabetically), (b) Steneck and Dethier’s classification (ordered by degree of cortication), (c) the binary canopy vs. turf and (d) the three-level canopy/subcanopy/turf classifications of vertical space use, and emergent groups created by *post hoc* clustering of our data using (e) agglomerative Hierarchical Clustering Analysis (HCA) and (f) the divisive *k*-medoids method. Within each grouping approach, stacked horizontal bars correspond to individual species. The order of individual species remains consistent across columns and is initially ordered according to Littler and Littler’s groups. Tracking individual species from left to right shows re-classification across grouping methods.

The emergent groups based on *post-hoc* clustering allow us to examine how species could be placed into five groups (matching the number of groups of the traditional schemes) to maximize explained variance. Most species were vastly reworked in their group allocation, with only a few tight nuclei of species left unchanged. (Thick) leathery species became split into two different clusters under both emergent grouping approaches (Figs 4, 5, Tables S2, S7). The first cluster, emergent group 2, corresponded to species that typically display high maximum length and SA:P. The second cluster, emergent group 3, mostly represented species with high C:N and thickness. Most species that belonged to the much spread out ‘coarsely branched’ and the corresponding ‘corticated’ groups were re-allocated across all emergent groups, leaving a tight nucleus of species unchanged (emergent group 1; Figs 4, 5, Tables S2, S7). Filamentous species tended to be grouped together regardless of the clustering approach (emergent group 4), corresponding mainly to high-SA:V species. The low degree of overlap between traditional and emergent functional groups underlines that current group allocations are far from optimal solutions to capture species’ dispersion in trait space.

## 4. DISCUSSION

As rapid global change dramatically alters community composition, it is imperative that functional schemes – across all major producer groups – are fit-for-purpose. We assessed whether common classifications of form and stature explained trait differences across macroalgae collected from rocky shores with, to our knowledge, the largest set of macroalgal traits assembled to date. Despite their convenience and prevalence in macroalgal ecology, the traditional classifications all left substantial amounts of interspecific trait variation unexplained. These results highlight the need for a re-evaluation of macroalgal grouping approaches and an increased focus on underlying functional trait variation.

### 4.1. Traditional groups poorly represent underlying physiological and ecological variation

Groups established by Littler and Littler (1980) and Steneck and Dethier (1994) were built on the premise that morphology and anatomy capture interspecific differences in physiology and function. Our finding of limited precision, accuracy, and stability of such groups suggests that anatomy (mainly, cortication) and morphology (gross form and branching order) fail to reliably capture variation in functional traits among species. The considerably better performance of emergent grouping is to be expected due to its *post hoc* nature, but indicates that the underlying limitation of traditional categorisations of macroalgal form is not grouping *per se*, but the very basis of groups in gross anatomy and morphology. Assuming functional traits are more direct proxies of physiology and potential contributions to ecosystem functions, Littler and Littler’s and Steneck and Dethier’s schemes do not provide a strong link between form and function.

In seeming contrast to our results, traditional classifications have been reported to account for functional properties, from photosynthetic rates to susceptibility to grazing (e.g., Littler & Littler, 1980, 1984; Steneck & Dethier, 1994). However, closer examination of these studies indicates that large overlap between groups is common and inferences are often drawn from differences between extreme groups, such as between jointed calcareous or encrusting against remaining groups (Littler & Arnold, 1982; Littler & Littler, 1984; Littler, Littler, & Taylor, 1983). Moreover, previous qualitative review found a general mismatch between functional groups and ecological responses and effects across ecosystems (Padilla & Allen, 2000). Our findings also echo work on vascular plants showing that growth form does not accurately represent underlying evolutionary tradeoffs between photosynthetic rates, construction costs, and longevity and hence, fails to capture global patterns of plant functional traits, nutrient cycling, and primary productivity (Díaz et al., 2016; Shipley et al., 2006).

Notwithstanding the overall inadequacy of traditional classifications of macroalgal form, some of their groups fared worse than others. The groups ‘(thick) leathery’ and ‘coarsely branched’/‘corticated’ were particularly spread out in trait space and were extensively reworked across grouping approaches, highlighting their lack of functional distinction. The ‘(thick) leathery’ group was split up between two main emergent clusters: an isolated one that included a few coarsely branched/corticated species, representing what may be considered ‘true’ leathery species, and one that associated with sheets. In fact, when using agglomerative clustering, the latter group consisted of high-surface area, sheet-like kelps (orders Laminariales and Tilopteridales) with a single leaf-life blade (e.g., *Alaria esculenta*) or digitate fronds (e.g., *Laminaria* spp.). ‘True’ leathery species were much more varied in morphology and mainly consisted of high-thickness, high-C:N fucophycids (subclass Fucophycidae, orders Fucales and Chordales; Algaebase, 2019). These species also had pneumatocysts, which seemed to be a strong determinant of group allocation among leathery macroalgae. Conversely, the filamentous group appeared to be the most stable across form-related traditional and emergent classifications. This suggests that filamentous species tend to display rather specific suites of traits that translate into precise anatomical and morphological features and hence, that are well-captured by the schemes of Littler and Littler and Steneck and Dethier. Such differences in the functional accuracy of form-related groups has implications for future applications of Littler and Littler’s and Steneck and Dethier’s schemes. We recommend that the ‘coarsely branched’/‘corticated’ group be more finely defined to reduce its breadth – perhaps requiring the creation of one or several additional groups guided by trait measurements – and that ‘(thick) leathery’ be separated into two groups to reflect the observed functional gap between ‘true’ (i.e., fucophycids) and sheet-like (i.e., kelps) leathery species.

### 4.2. Common classifications of vertical space use provide coarse assessments of the ecological impacts of fast-paced changes in macroalgal assemblages

Classifications of vertical space use are widely used because of their convenience in accounting for changes in macroalgal community composition (Filbee-Dexter & Wernberg, 2018). These groups were the least parsimonious of approaches, accounted for little multivariate trait expression, and showed a high degree of overlap for individual traits. Accordingly, turnover in species within these groups – or canopy to turf transitions – will have different ecological consequences depending on species traits. Recent focused comparative work on two kelp (canopy) species supports this point: along the southern coast of Britain under ocean warming, replacement of *Laminaria hyperborea* with *L. ochroleuca* results in ecologically significant changes in supported biodiversity and ecosystem processes (Pessarrodona et al., 2019; Smale, Wernberg, Yunnie, & Vance, 2015). Moreover, allocation of macroalgal species to categories of vertical space use is community-dependent and can be subjective, and various definitions of turfs have been employed, resulting in a lack of generality of those schemes across ecosystems (Connell, Foster, & Airoldi, 2014). Our findings imply that coarse categorisations of macroalgal stature will hinder explanatory power and the generality of global synthesis when assessing the ecological impacts of environment-mediated changes in macroalgal assemblages.

Nevertheless, classifications of macroalgal stature can remain insightful when applied to ecosystem functioning. Perhaps aided by greater within-group sample sizes, both classifications of vertical space use explained significant differences for most individual functional traits. The trait differences entailed in canopy-to-turf shifts, such as reduced C:N and higher STA, correspond to contrasts between ‘slower’, longer-lived species, and ‘faster’ opportunistic forms (Airoldi, 1998; Littler & Littler, 1980, 1981). These differences may partially explain the global rise of turfs with altered environmental conditions (Feehan et al., 2019; Filbee-Dexter & Wernberg, 2018) as well as the consequences of canopy-to-turf shifts for carbon flow and habitat provisioning (Copertino, Connell, & Cheshire, 2005; Filbee-Dexter & Wernberg, 2018). Therefore, despite risking substantial loss of information and lack of generality, broad classifications of macroalgal stature may offer quick first-order assessments of the ecological impacts of fast-paced changes in macroalgal assemblages.

### 4.3 A call for global trait-based functional ecology in macroalgae

Macroalgal traits – and by extension the responses of macroalgae to environmental change and the consequences for ecosystem function – are poorly captured by traditional grouping approaches. We call for a standardized trait-based approach to macroalgal ecology in order to more accurately and generally quantify their ecological differences. The suite of traits measured here is by no means exhaustive, but illustrates the potential for standardised screening of traits in macroalgae, despite their extensive evolutionary history and morphological diversity. Given the labour intensity of trait screening, a key challenge is to identify a small set of – ideally orthogonal – ecologically relevant traits before embarking on globally coordinated efforts. Here, maximum length and branching order emerge as contenders because they were most associated with orthogonal axes of variation and have the benefit of being relatively easy to measure. Ultimately and similarly to plant functional types, emergent groups, if found to be consistent across species and environmental conditions, have the potential to represent macroalgal variation more closely than traditional groups and to provide prompt assessments of the ecological impacts of macroalgal community change.

## 5. CONCLUSION

We found little support for the group gambit of macroalgae: gross morphology, anatomy and relative stature appear to inadequately capture functional trait variation. Trait-based approaches have the potential to represent macroalgal functional variation more closely and therefore to provide stronger assessments of macroalgal beds and forests under global change. Studies like ours, which emphasise the potential of direct use of functional traits in capturing macroalgal form and function, can act as an incentive to build large, coordinated datasets spanning all main aspects of macroalgal eco-physiology. Ultimately, macroalgae could be placed along an extended spectrum of primary producer functional variation.

## ACKNOWLEDGEMENTS

This project was funded by a Marie Curie Integration Career grant to J. Griffin (number: FP7 MC CIG 61893). A. Mauffrey was also supported by a Swansea University research scholarship. We would like to thank A. Raduan, S. Ball, J. Hunt, P. Deacon, O. Smith, E. Milne, G. Burton, M. Georgiev, O. Koppel, L. Wójcik, J. Mutter, F. Guillen Ezcurra, G. Blow, D. de Battisti, and T. Fairchild, for the invaluable help with sampling and trait screening. We thank M. Wilkinson for his help during the 2017 British Phycological Society Seaweed Field Meeting in Orkney.

## AUTHORS’ CONTRIBUTIONS

AM and JG conceived the ideas, designed methodology, and led the writing of the manuscript; AM and LC collected the data and created the figures and tables; AM analysed the data with the help of JG and LC. All authors contributed critically to the drafts and gave final approval for publication.

## DATA ACCESSIBILITY STATEMENT

We intend to archive our data at Figshare.

## REFERENCES

Airoldi, L. (1998). Roles of Disturbance, Sediment Stress, and Substratum Retention on Spatial Dominance in Algal Turf. Ecology, 79(8), 2759–2770. doi: 10.1890/0012-9658(1998)079[2759:RODSSA]2.0.CO;2

Arenas, F., Sánchez, I., Hawkins, S. J., & Jenkins, S. R. (2006). The invasibility of marine algal assemblages: role of functional diversity and identity. Ecology, 87(11), 2851–2861.

Cappelatti, L., Mauffrey, A. R. L., & Griffin, J. N. (2019). Applying continuous functional traits to large brown macroalgae: variation across tidal emersion and wave exposure gradients. Marine Biology, 166(11), 145. doi: 10.1007/s00227-019-3574-5

Carpenter, R. C. (1990). Competition Among Marine Macroalgae: A Physiological Perspective. Journal of Phycology, 26(1), 6–12. doi: 10.1111/j.0022-3646.1990.00006.x

Chapin, F. S., Bret-Harte, M. S., Hobbie, S. E., & Zhong, H. (1996). Plant functional types as predictors of transient responses of arctic vegetation to global change. Journal of Vegetation Science, 7(3), 347–358. doi: 10.2307/3236278

Chung, K. I., Beardall, J., Mehta, S., Sahoo, D., & Stojkovic, S. (2011). Using marine macroalgae for carbon sequestration: a critical appraisal. Journal of Applied Phycology, 23(5), 877–886. doi: 10.1007/s10811-010-9604-9

Connell, S. D., Foster, M. S., & Airoldi, L. (2014). What are algal turfs? Towards a better description of turfs. Marine Ecology Progress Series, 495, 299–307. doi: 10.3354/meps10513

Copertino, M., Connell, S. D., & Cheshire, A. (2005). The prevalence and production of turf-forming algae on a temperate subtidal coast. Phycologia, 44(3), 241–248. doi: 10.2216/0031-8884(2005)44[241:TPAPOT]2.0.CO;2

Cornelissen, J. H. C., Lavorel, S., Garnier, E., Díaz, S., Buchmann, N., Gurvich, D. E., … Poorter, H. (2003). A handbook of protocols for standardised and easy measurement of plant functional traits worldwide. Australian Journal of Botany, 51, 335–380.

de los Santos, C. B., Onoda, Y., Vergara, J. J., Pérez-Lloréns, J. L., Bouma, T. J., Nafie, Y. A. L., … Brun, F. G. (2016). A comprehensive analysis of mechanical and morphological traits in temperate and tropical seagrass species. Marine Ecology Progress Series, 551, 81–94. doi: 10.3354/meps11717

Díaz, S., & Cabido, M. (2001). Vive la différence: plant functional diversity matters to ecosystem processes. Trends in Ecology and Evolution, 16, 646–655.

Díaz, S., Kattge, J., Cornelissen, J. H. C., Wright, I. J., Lavorel, S., Dray, S., … Gorné, L. D. (2016). The global spectrum of plant form and function. Nature, 529(7585), 167–171. doi: 10.1038/nature16489

Díaz, S., Purvis, A., Cornelissen, J. H. C., Mace, G. M., Donoghue, M. J., Ewers, R. M., … Pearse, W. D. (2013). Functional traits, the phylogeny of function, and ecosystem service vulnerability. Ecology and Evolution, 3(9), 2958–2975. doi: 10.1002/ece3.601

Dromgoole, F. I. (1981). Form and Function of the Pneumatocysts of Marine Algae. I. Variations in the Pressure and Composition of Internal Gases. Botanica Marina, 24, 257–266. doi: 10.1515/botm.1981.24.5.257

Elger, A., & Willby, N. J. (2003). Leaf dry matter content as an integrative expression of plant palatability: the case of freshwater macrophytes. Functional Ecology, 17, 58–65.

Feehan, C. J., Grace, S. P., & Narvaez, C. A. (2019). Ecological feedbacks stabilize a turf-dominated ecosystem at the southern extent of kelp forests in the Northwest Atlantic. Scientific Reports, 9(1), 1–10. doi: 10.1038/s41598-019-43536-5

Filbee-Dexter, K., & Wernberg, T. (2018). Rise of Turfs: A New Battlefront for Globally Declining Kelp Forests. BioScience, 68(2), 64–76. doi: 10.1093/biosci/bix147

Garnier, E., Cortez, J., Billès, G., Navas, M.-L., Roumet, C., Debussche, M., … Toussaint, J.-P. (2004). Plant functional markers capture ecosystem properties during secondary succession. Ecology, 85(9), 2630–2637.

Google Scholar. (2019). Retrieved October 10, 2019, from Google Scholar UK: https://scholar.google.co.uk/

Griffiths, M., Harrison, S. T. L., Smit, M., & Maharajh, D. (2016). Major Commercial Products from Micro- and Macroalgae. In F. Bux & Y. Chisti (Eds.), Algae Biotechnology: Products and Processes (pp. 269–300). doi: 10.1007/978-3-319-12334-9_14

Guiry, M., & Guiry, G. (2019). Algaebase. Retrieved October 10, 2019, from Algaebase website: https://www.algaebase.org

Hanisak, M. D., Littler, M. M., & Littler, D. S. (1988). Significance of macroalgal polymorphism: intraspecific tests of the functional-form model. Marine Biology, 99, 157–165.

Harley, C. D. G., Anderson, K. M., Demes, K. W., Jorve, J. P., Kordas, R. L., Coyle, T. A., & Graham, M. H. (2012). Effects of Climate Change on Global Seaweed Communities. Journal of Phycology, 48(5), 1064–1078. doi: 10.1111/j.1529-8817.2012.01224.x

Hay, M. E. (1981). The functional morphology of turf forming seaweeds: persistence in stressful marine habitats. Ecology, 62, 739–750.

Hurd, C. L., Harrison, P. J., Bischof, K., & Lobban, C. S. (2014). Seaweed Ecology and Physiology. Cambridge University Press.

Kassambara, A. (2019). ggpubr: “ggplot2” Based Publication Ready Plots (Version 0.2.1) [R, R (≥ 3.1.0), ggplot2, magrittr]. Retrieved October 10, 2019, from https://cloud.r-project.org/web/packages/ggpubr/index.html

Lavorel, S., & Garnier, E. (2002). Predicting changes in community composition and ecosystem functioning from plant traits: revisiting the Holy Grail. Functional Ecology, 16, 545–556.

Lavorel, S., McIntyre, S., Landsberg, J., & Forbes, T. D. A. (1997). Plant functional classifications: from general groups to specific groups based on response to disturbance. Trends in Ecology and Evolution, 12, 474–478.

Littler, M. M. (1980). Morphological form and photosynthesis performances of marine macroalgae: tests of a functional/form hypothesis. Botanica Marina, 22, 161–165.

Littler, M. M., & Arnold, K. E. (1982). Primary productivity of marine macroalgal functional-form groups from Southwestern North America. Journal of Phycology, 18, 307–311.

Littler, M. M., & Littler, D. S. (1980). The evolution of thallus form and survival strategies in benthic marine macroalgae: field and laboratory tests of a functional form model. American Naturalist, 116(1), 25–44.

Littler, M. M., & Littler, D. S. (1981). Intertidal macrophyte communities from Pacific Baja California and the upper gulf of California: relatively constant vs. environmentally fluctuating systems. Marine Ecology Progress Series, 4, 145–158.

Littler, M. M., & Littler, D. S. (1984). Relationships between macroalgal functional form groups and substrata stability in a subtropical rocky-intertidal system. Journal of Experimental Marine Biology and Ecology, 74(1), 13–34.

Littler, M. M., Littler, D. S., & Taylor, P. R. (1983). Evolutionary strategies in a tropical barrier reef system: functional-form groups of marine macroalgae. Journal of Phycology, 19, 229–237.

Littler, M. M., Taylor, P. R., & Littler, D. S. (1983). Algal resistance to herbivory on a Caribbean barrier reef. Coral Reefs, 2(2), 111–118. doi: 10.1007/BF02395281

Lukasová, A. (1979). Hierarchical agglomerative clustering procedure. Pattern Recognition, 11(5-6), 365–381. doi: 10.1016/0031-3203(79)90049-9

Maechler, M., Rousseeuw, P., Struyf, A., Hubert, M., & Hornik, K. (2019). cluster: Cluster Analysis Basics and Extensions (Version 2.1.0) [R, R (≥ 3.3.0)]. Retrieved October 10, 2019, from https://cran.r-project.org/web/packages/cluster/index.html

Mazel, F., Pennell, M. W., Cadotte, M. W., Diaz, S., Riva, G. V. D., Grenyer, R., … Pearse, W. D. (2018). Prioritizing phylogenetic diversity captures functional diversity unreliably. Nature Communications, 9(1), 1–9. doi: 10.1038/s41467-018-05126-3

McGill, B. J., Enquist, B. J., Weiher, E., & Westoby, M. (2006). Rebuilding community ecology from functional traits. Trends in Ecology and Evolution, 21(4), 178–185.

McLachlan, J. (1985). Macroalgae (seaweeds): industrial resources and their utilization. Plant and Soil, 89, 137–157.

Miranda, M., Lopez-Alonso, M., & Garcia-Vaquero, M. (2017). Macroalgae for Functional Feed Development: Applications in Aquaculture, Ruminant and Swine Feed Industries. In Veterinary Medicine Research Collection. Seaweeds: Biodiversity, Environmental Chemistry and Ecological Impacts (NOVA science publishers). Retrieved October 10, 2019, from https://researchrepository.ucd.ie/handle/10197/9045

Murtagh, F., & Legendre, P. (2014). Ward’s Hierarchical Agglomerative Clustering Method: Which Algorithms Implement Ward’s Criterion? Journal of Classification, 31(3), 274–295. doi: 10.1007/s00357-014-9161-z

Oksanen, J., Blanchet, F. G., Friendly, M., Kindt, R., Legendre, P., McGlinn, D., … Wagner, H. (2019). vegan: Community Ecology Package (Version 2.5-5) [R, R (≥ 3.4.0), permute (≥ 0.9-0), lattice]. Retrieved October 10, 2019, from https://cran.r-project.org/web/packages/vegan/index.html

Padilla, D. K. (1984). The importance of form: differences in competitive ability, resistance to consumers and environmental stress in an assemblage of coralline algae. Journal of Experimental Marine Biology and Ecology, 79, 105–127.

Padilla, D. K., & Allen, B. J. (2000). Paradigm lost: reconsidering functional form and group hypotheses in marine ecology. Journal of Experimental Marine Biology and Ecology, 250, 207–221.

Pessarrodona, A., Foggo, A., & Smale, D. A. (2019). Can ecosystem functioning be maintained despite climate-driven shifts in species composition? Insights from novel marine forests. Journal of Ecology, 107(1), 91–104. doi: 10.1111/1365-2745.13053

Petchey, O. L., & Gaston, K. J. (2006). Functional diversity: back to basics and looking forward. Ecology Letters, 9(6), 741–758. doi: 10.1111/j.1461-0248.2006.00924.x

Phillips, J. C., Kendrick, G. A., & Lavery, P. S. (1997). A test of a functional group approach to detecting shifts in macroalgal communities along a disturbance gradient. Marine Ecology Progress Series, 153, 125–138.

Pianka, E. R. (1970). On *r*- and *K*-Selection. The American Naturalist, 104(940), 592–597. doi: 10.1086/282697

R Core Team. (2019). R: A Language and Environment for Statistical Computing (Version 3.5.3) [R]. Retrieved October 10, 2019, from http://www.R-project.org/.

Reich, P. B. (2014). The world-wide ‘fast–slow’ plant economics spectrum: a traits manifesto. Journal of Ecology, 102(2), 275–301. doi: 10.1111/1365-2745.12211

Reynolds, A. P., Richards, G., de la Iglesia, B., & Rayward-Smith, V. J. (2006). Clustering Rules: A Comparison of Partitioning and Hierarchical Clustering Algorithms. Journal of Mathematical Modelling and Algorithms, 5(4), 475–504. doi: 10.1007/s10852-005-9022-1

Roos, R. E., Zuijlen, K. van, Birkemoe, T., Klanderud, K., Lang, S. I., Bokhorst, S., … Asplund, J. (in press). Contrasting drivers of community-level trait variation for vascular plants, lichens, and bryophytes across an elevational gradient. Functional Ecology. doi: 10.1111/1365-2435.13454

Schneider, C. A., Rasband, W. S., & Eliceiri, K. W. (2012). NIH Image to ImageJ: 25 years of image analysis. Nature Methods, 9(7), 671–675. doi: 10.1038/nmeth.2089

Schonbeck, M. W., & Norton, T. A. (1979). Drought-Hardening in the Upper-Shore Seaweeds *Fucus spiralis* and *Pelvetia canaliculata*. Journal of Ecology, 67(2), 687–696. doi: 10.2307/2259120

Shipley, B., Lechowicz, M. J., Wright, I., & Reich, P. B. (2006). Fundamental trade-offs generating the worldwide leaf economics spectrum. Ecology, 87(3), 535–514.

Smale, D. A., Wernberg, T., Yunnie, A. L. E., & Vance, T. (2015). The rise of *Laminaria ochroleuca* in the Western English Channel (UK) and comparisons with its competitor and assemblage dominant *Laminaria hyperborea*. Marine Ecology, 36(4), 1033–1044. doi: 10.1111/maec.12199

Smart, S. M., Glanville, H. C., Blanes, M. del C., Mercado, L. M., Emmett, B. A., Jones, D. L., … Hodgson, J. G. (2017). Leaf dry matter content is better at predicting above-ground net primary production than specific leaf area. Functional Ecology, 1336–1344. doi: 10.1111/1365-2435.12832

Starko, S., Claman, B. Z., & Martone, P. T. (2015). Biomechanical consequences of branching in flexible wave-swept macroalgae. New Phytologist, 206(1), 133–140. doi: 10.1111/nph.13182

Steneck, R. S., & Dethier, M. N. (1994). A functional group approach to the structure of algal-dominated communities. Oikos, 69, 476–498.

Stewart, H. L., & Carpenter, R. C. (2003). The effects of morphology and water flow on photosynthesis of marine macroalgae. Ecology, 84(11), 2999–3012.

Suding, K. N., Lavorel, S., Chapin, F. S. I., Cornelissen, J. H. C., Díaz, S., Garnier, E., … Navas, M.-L. (2008). Scaling environmental change through the community-level: a trait-based response-and-effect framework for plants. Global Change Biology, 14, 1125–1140. doi: 10.1111/j.1365-2486.2008.01557.x

Taylor, P. R., & Hay, M. E. (1984). Functional morphology of intertidal seaweeds: adaptive significance of aggregate vs. solitary forms. Marine Ecology Progress Series, 18, 295–302.

Teagle, H., Hawkins, S. J., Moore, P. J., & Smale, D. A. (2017). The role of kelp species as biogenic habitat formers in coastal marine ecosystems. Journal of Experimental Marine Biology and Ecology, 492, 81–98. doi: 10.1016/j.jembe.2017.01.017

Trugman, A. T., Anderegg, L. D. L., Wolfe, B. T., Birami, B., Ruehr, N. K., Detto, M., … Anderegg, W. R. L. (2019). Climate and plant trait strategies determine tree carbon allocation to leaves and mediate future forest productivity. Global Change Biology, 25(10), 3395–3405. doi: 0.1111/gcb.14680

van Buuren, S., & Groothuis-Oudshoorn, K. (2011). mice: Multivariate Imputation by Chained Equations. Journal of Statistical Software, 45(3), 1–67.

Veiga, P., Rubal, M., & Sousa-Pinto, I. (2014). Structural complexity of macroalgae influences epifaunal assemblages associated with native and invasive species. Marine Environmental Research, 101, 115–123.

Violle, C., Enquist, B. J., McGill, B. J., Jiang, L., Albert, C. H., Hulshof, C., … Messier, J. (2012). The return of the variance: intraspecific variability in community ecology. Trends in Ecology and Evolution, 27(4), 244–252. doi: 10.1016/j.tree.2011.11.014

Weykam, G., Gómez, I., Wiencke, C., Iken, K., & Klöser, H. (1996). Photosynthetic characteristics and C:N ratios of macroalgae from King George Island (Antarctica). Journal of Experimental Marine Biology and Ecology, 204, 1–22.

Wickham, H. (2009). ggplot2: Elegant Graphics for Data Analysis (2nd ed., Vols. 1-1). Springer International Publishing.

Wilson, P., Thompson, K., & Hodgson, J. G. (1999). Specific leaf area and leaf dry matter content as alternative predictors of plant strategies. New Phytologist, 143, 155–162.

Wright, I. J., Reich, P. B., Westoby, M., Ackerly, D. D., Baruch, Z., Bongers, F., … Villar, R. (2004). The worldwide leaf economics spectrum. Nature, 428(6985), 821–827. doi: 10.1038/nature02403

