## Supporting information for "Traditional functional groups capture limited variation in the trait space of macroalgae"

**APPENDIX**

1. **Steneck and Dethier’s detailed classification provides mostly functionally-redundant information**

**Introduction**

Steneck and Dethier (1994) invited macroalgal ecologists to use a finer scale of morphological (mainly, branching pattern) and anatomical (mainly, degree of cortication) divisions than their five-group classification. Two of the main groups, ‘filamentous (S)’ and ‘foliose’, can be further subdivided into the sub-groups ‘corticated filamentous’ for thinly corticated and/or polysiphonious species and ‘corticated foliose’ for sheet species possessing a distinct layer of cortical cells.

**Methods**

We additionally ran all analyses using seven clusters to match Steneck and Dethier’s detailed classification and to assess the influence of increasing the number of functional groups on explaining macroalgal functional variability. Mirroring five-group clustering, we created emergent groups from the weighted Gower matrix using seven-cluster agglomerative HCA and *k*-medoids. We ran three additional PERMANOVAs to those described in the main text on the weighted Gower matrix we created, using Steneck and Dethier’s seven-group classification and the two seven-cluster emergent classifications as a grouping variable. We ran the PERMANOVAs on all traits combined as well as solely on traits known to relate to economics and competitive dominance, so as to assess the extent of trait variation explained along the two major axes of ecological variability found amongst global plant species (Díaz et al., 2016). We assessed the parsimony of the PERMANOVAs using Akaike’s Information Criterion.

**Results**

Similarly to Steneck and Dethier’s five-cluster classification, their detailed, seven-cluster scheme showed extensive group overlap across traits, with only 46 % of all possible pairwise group pairs being significantly different (Figs S9, S10, Table S8). The detailed classification explained ca. 41 % of multivariate trait variation (seven-cluster PERMANOVA, *R*² = 0.4143, *P* < 0.001). Increasing the number of clusters from five to seven while using Steneck and Dethier’s classification was linked to about 21 % of species being re-allocated to different groups and explained about 4 % more multivariate trait variation with greater parsimony (AIC = 60.75 for Steneck and Dethier’s seven-group classification against 63.63; Fig. S10, S11, Table S7). However, most of the extra trait variation accounted for by Steneck and Dethier’s seven-cluster classification was functionally redundant with its five-cluster counterpart. Subgroups of the seven-cluster classification were located within their hierarchical superiors (groups 2 and 3), creating a bullseye pattern on the biplots (Fig. S10, Table S2).

Seven-cluster emergent groups explained ca. 64 (seven-cluster PERMANOVA, *R*² = 0.6428, *P* < 0.001) and 63 % (seven-cluster PERMANOVA, *R*² = 0.6332, *P* < 0.001) of multivariate trait variation using agglomerative HCA and the divisive *k*-medoids analysis, respectively. This corresponds to an increase of about 9 % of trait variation explained when compared to the classifications created by five-cluster HCA and *k*-medoids. Both seven-cluster emergent classifications showed the greatest parsimony of all grouping approaches, with HCA leading (AIC = 13.77 and 16.28 for HCA and *k*-medoids, respectively). While the two seven-cluster emergent classifications explained just about the same amount of trait variation, ca. 23 % of the species were re-allocated to different groups across both approaches (Fig. S10, S11, Table S7). Groups that were most affected by changing the cluster number from five to seven were those that were analogous to (thick) leathery and sheet/foliose and their reworked emergent group analogues (groups 2 and 3); they were further split up (Figs 3, 4, S11, S12, Table S7).

**Discussion**

Steneck and Dethier’s (1994) complete classification explained little more multivariate trait variation than its simplified, five-cluster counterpart. Most of the extra multivariate trait variation captured was simply due to the greater number of divisions, which automatically helps explain greater variation, while the additional groups captured mostly redundant information in trait space. Overall, such results confirm that traditional grouping approaches do not accurately represent underlying functional trait variability and in fact, have little basis from a functional standpoint. Our study suggests that the low explanatory power of traditional grouping approaches stems from both the low group number and functional inadequacy of the groups themselves (particularly ‘coarsely branched’/‘corticated’ and ‘(thick) leathery’). When using emergent groups though, adding two additional groups to reach a total of seven clusters explained substantially more trait variation in a more parsimonious way, which confirms that beside their functional inadequacy, traditional grouping approaches are also likely to be constrained by the low number of their groups and hence, their coarseness. Increasing the number of groups to at least seven in future morphology- and anatomy-based or emergent classifications may offer greater precision in capturing underlying functional trait variability.

2. **List of tables**

Table S1. Functional traits measured and their physiological significance.

Table S2. Species allocation to traditional and emergent groups.

Table S3. Site details.

Table S4. Sampling and trait screening methodology.

Table S5. Trait transformations and weightings used in the Gower matrix.

Table S6. Spearman's rank correlations between the functional traits studied.

Table S7. Differences in species composition across seven-cluster grouping methods (%).

3. **List of figures**

Figure S1. Intra- and interspecific variability among the continuous functional traits measured.

Figure S2. Distribution of functional groups along the first two principal component axes.

Figure S3. Distribution of species-level traits among Steneck and Dethier’s groups.

Figure S4. Pairwise differences among Littler and Littler’s groups.

Figure S5. Pairwise differences among Steneck and Dethier’s groups.

Figure S6. Distribution of species-level traits among a common three-level classification of vertical space use.

Figure S7. Pairwise differences among the groups of a common three-level classification of macroalgal vertical space use.

Figure S8. Distribution of species-level traits among the groups established by Steneck and Dethier’s seven-cluster classification.

Figure S9. Pairwise differences among the groups of Steneck and Dethier’s seven-cluster classification.

Figure S10. Distribution of seven-cluster functional groups along the first two principal coordinate axes.

Figure S11. Species re-allocation across seven-cluster grouping approaches.

**Table S1. Functional traits and their physiological significance.** A question mark indicates that the functional trait and either spectrum of ecological variation, economics or competitive dominance, may not relate in a straightforward way. References are given in superscript and are the following: 1. Littler and Littler (1980); 2. Dromgoole (1981); 3. Hay (1981); 4. Taylor and Hay (1984); 5. Carpenter (1990); 6. Weykam et al. (1996); 7. Reich et al. (1999); 8. Roderick et al. (2000); 9. Cornelissen et al. (2003); 10. Elger and Willby (2003); 11. Vile et al. (2005); 12.Veiga et al. (2014).

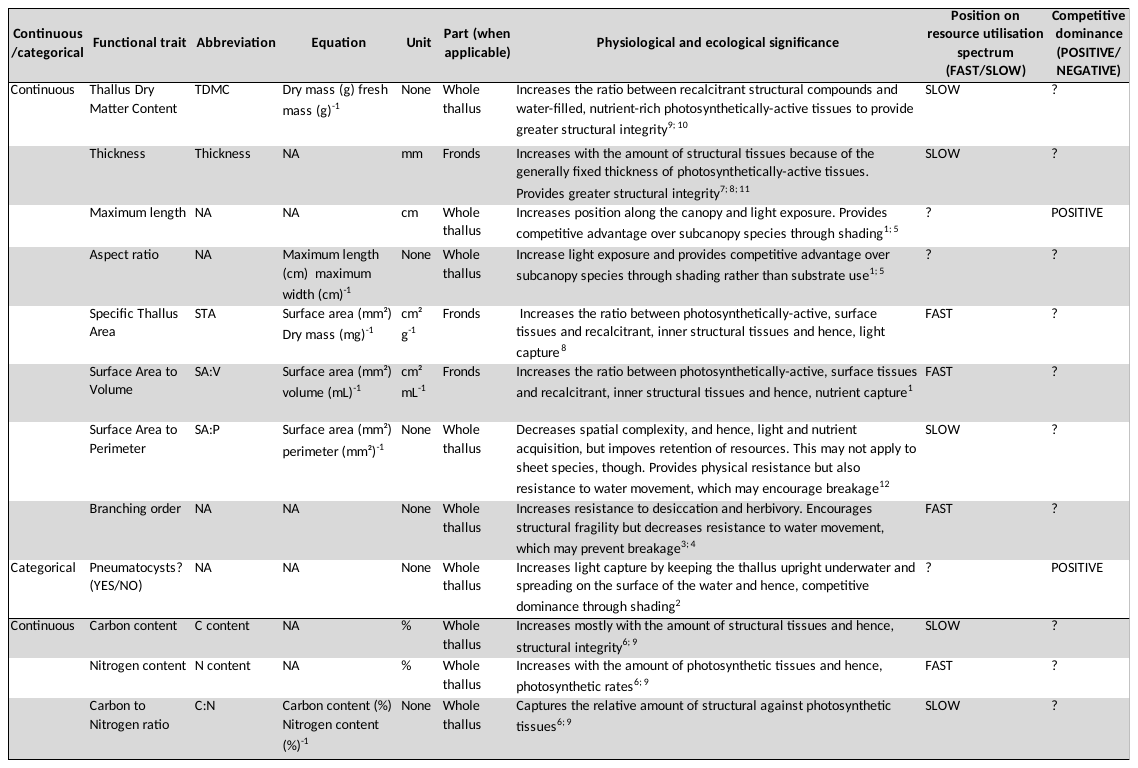

**Table S2. Species allocation to functional groups.** Species were attributed to Littler and Littler's (1980) and Steneck and Dethier's (1994) groups after a review of the literature, and to groups relating to vertical space use (canopy vs. turf and canopy/subcanopy/turf) based on observed patterns at the sites screened. We also indicate species allocation across five- and seven-cluster emergent classifications created by *post hoc* clustering of multivariate trait values using agglomerative Hierarchical Clustering Analysis (HCA) and the divisive *k*-medoids method (*k*). References given in superscript are the following: 1. Littler (1980); 2. Littler and Littler (1980); 3. Littler and Arnold (1982); 4. Littler and Taylor (1983); 5. Littler et al. (1983); 6. Littler and Littler (1984); 7. Steneck and Dethier (1994); 8. Sánchez and Fernández (2006); 9. Bates (2009); 10. Gómez and Huovinen (2011); 11. Hurd et al. (2014).

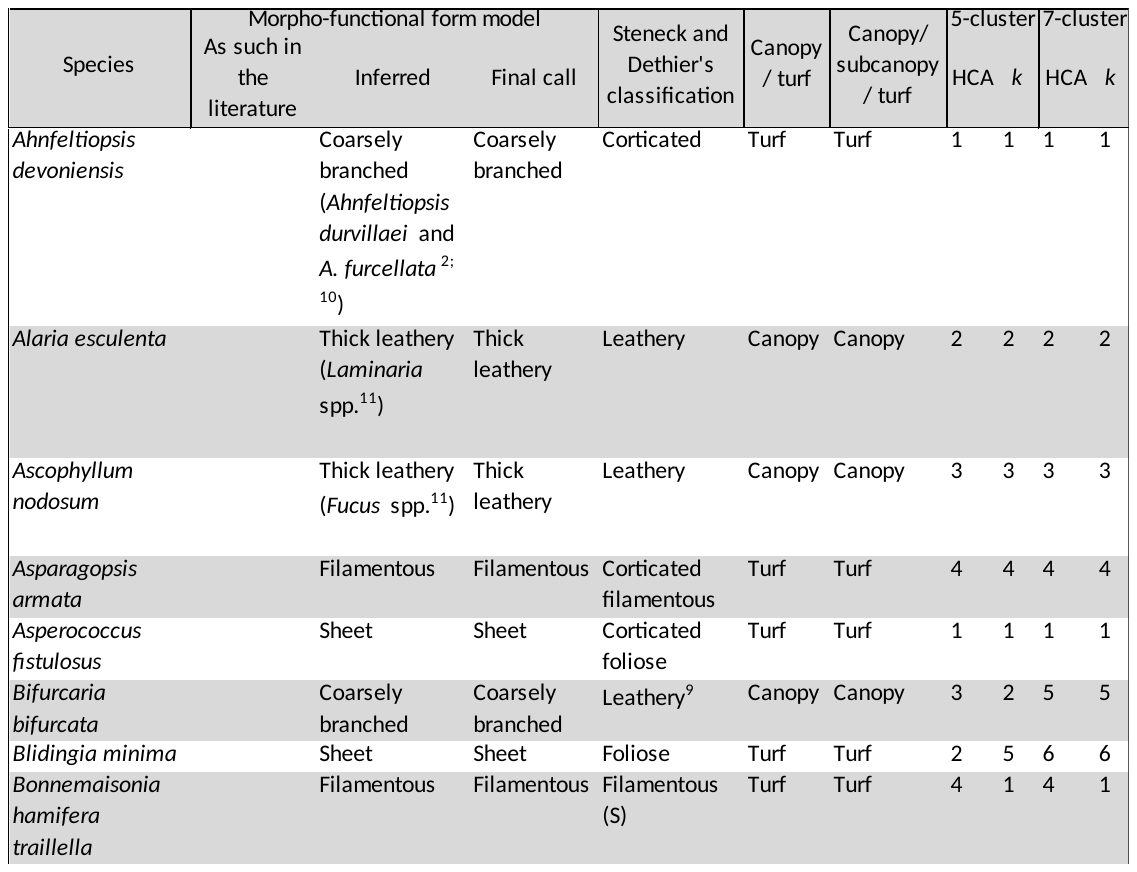

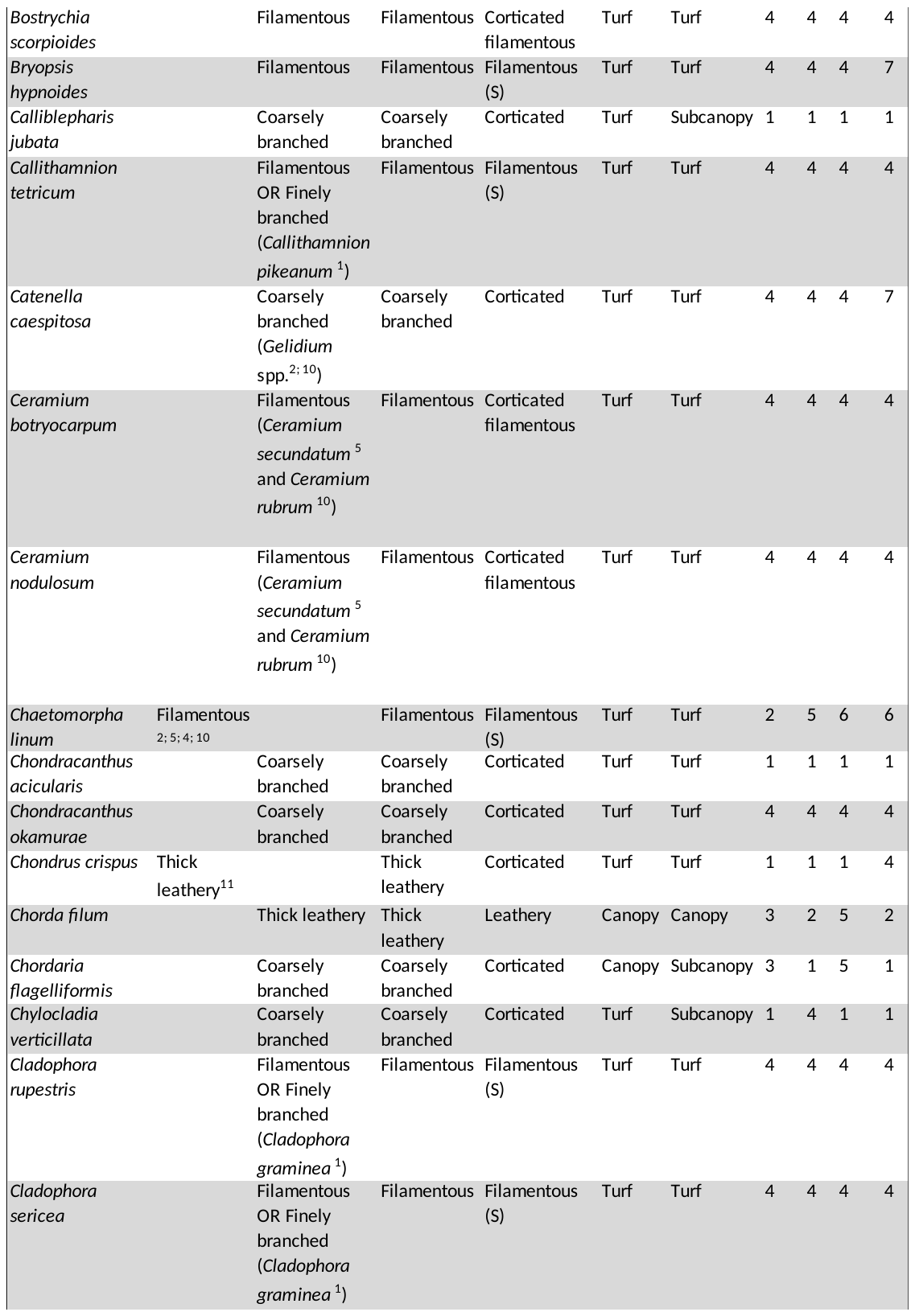

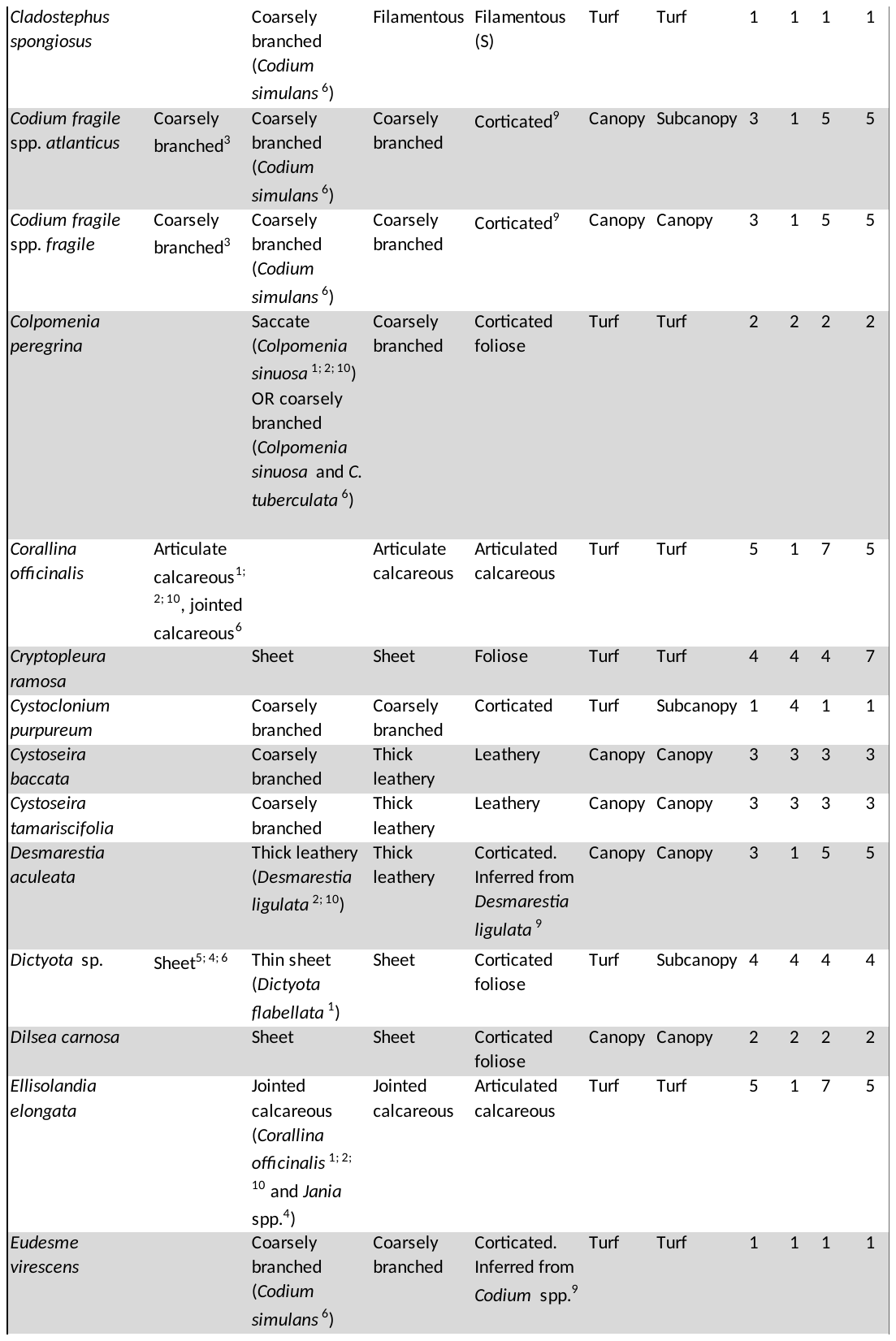

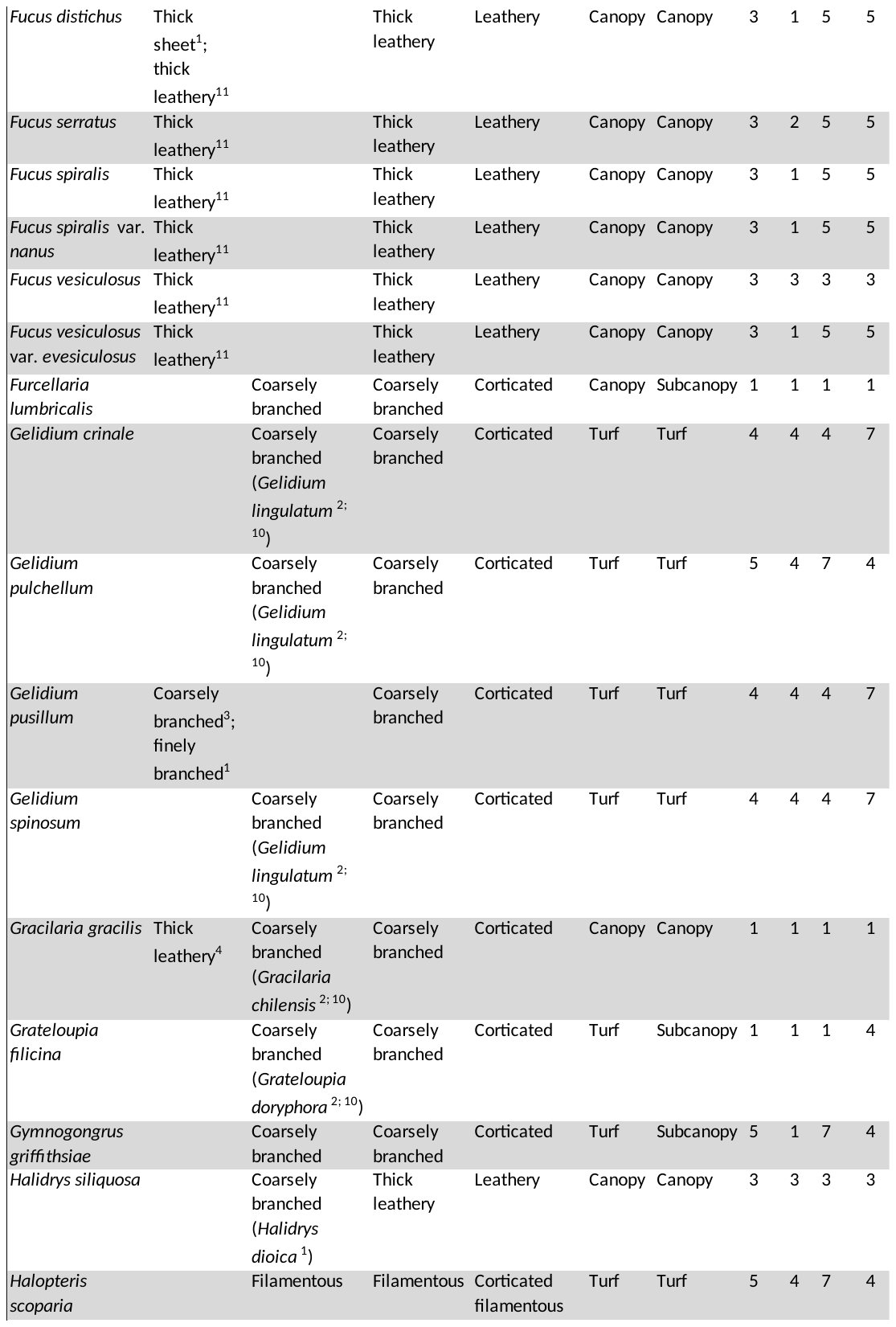

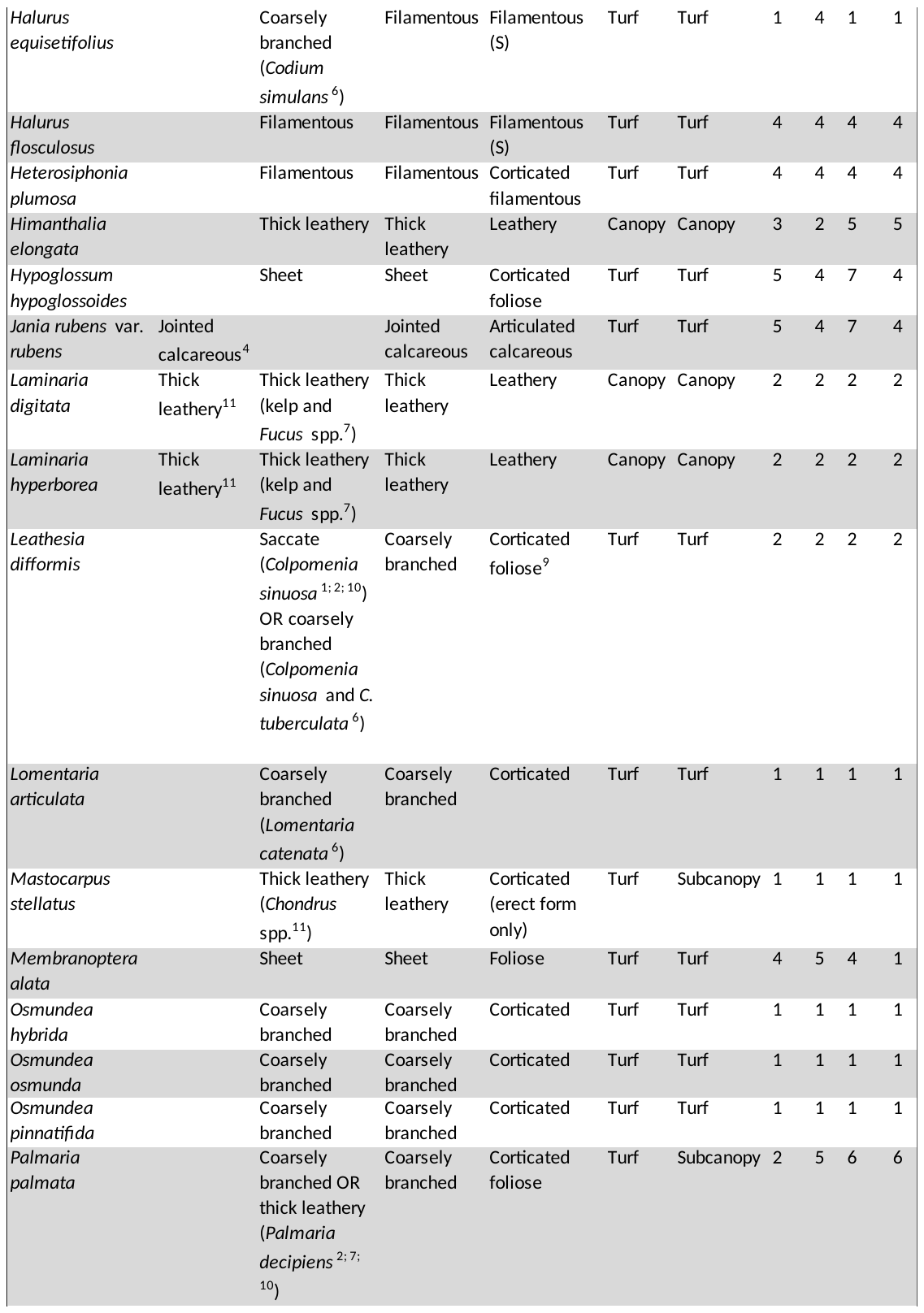

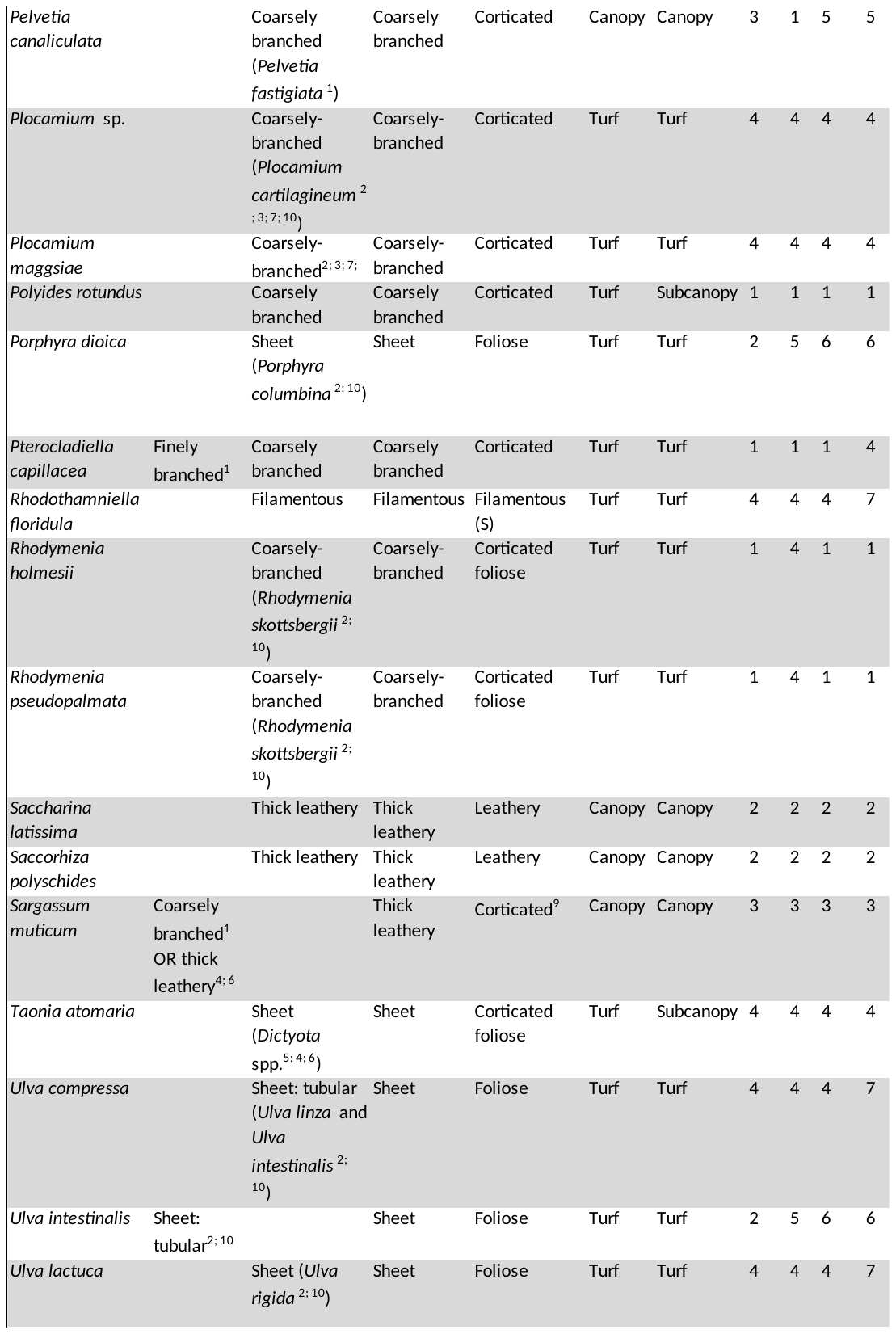

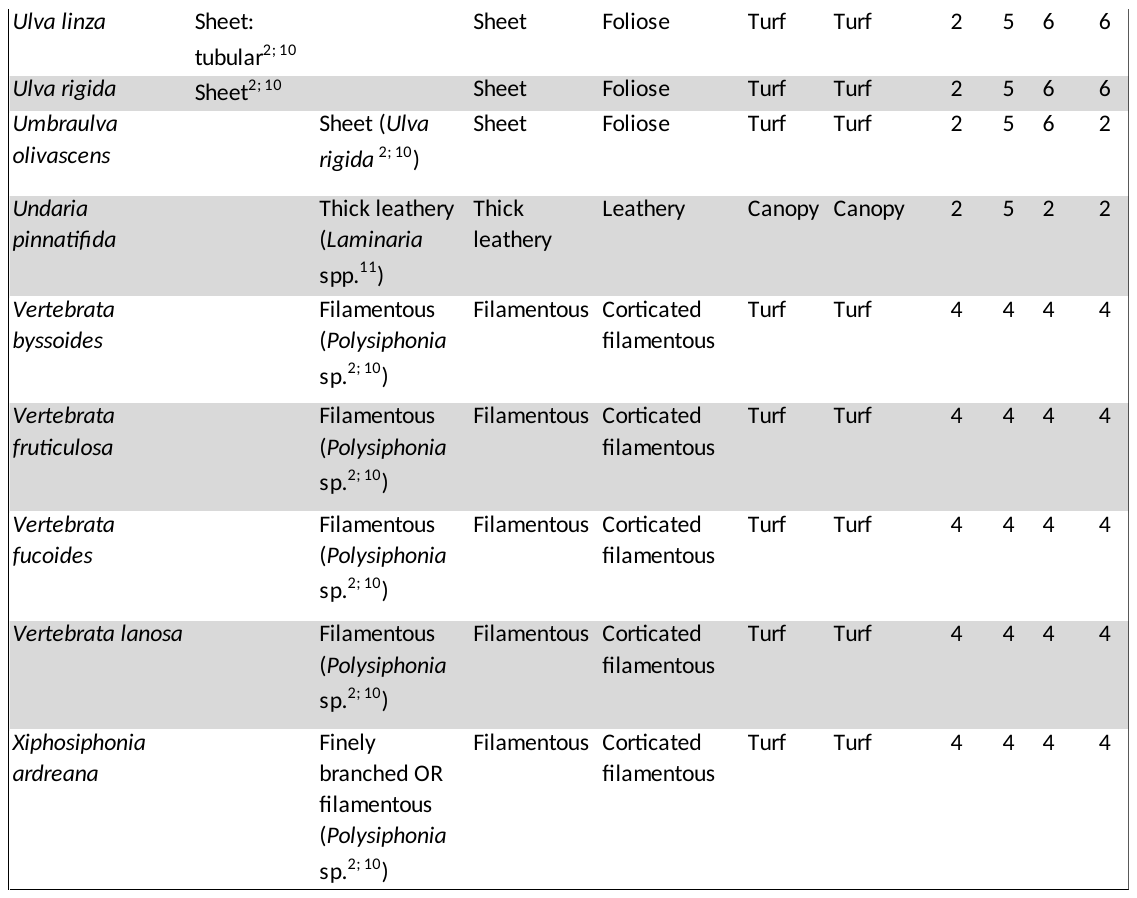

**Table S3. Site details.** We collected samples from twelve intertidal rocky shores in the UK ranging from very sheltered to very exposed.

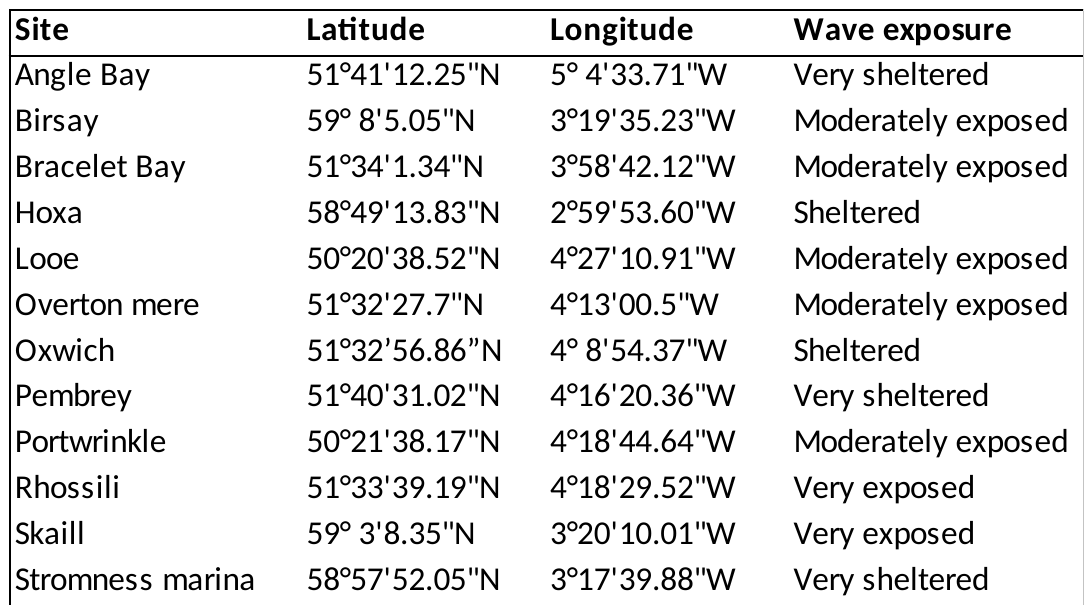

**Table S4. Sampling and trait screening methodology.** Year and month of collection, site, and number of replicates (reps.) per species are shown for the 95 intertidal macroalgal species screened. Site names are abbreviated as follows: ‘A’ for Angle Bay, ‘B’ for Birsay, ‘BB’ for Bracelet Bay, ‘H’ for Hoxa, ‘L’ for Looe, ‘O’ for Overton mere, ‘Ox’ for Oxwich, ‘P’ for Pembrey, ‘Pw’ for Portwrinkle, ‘R’ for Rhossili, ‘S’ for Skaill, and ‘SM’ for Stromness marina (Table S3). We also indicate whether the samples were made up of a single or multiple individual(s; ind.) or tufts (when turf-forming), processed fresh or frozen, subsampled, and microscopically photographed for surface-area and perimeter measurements.

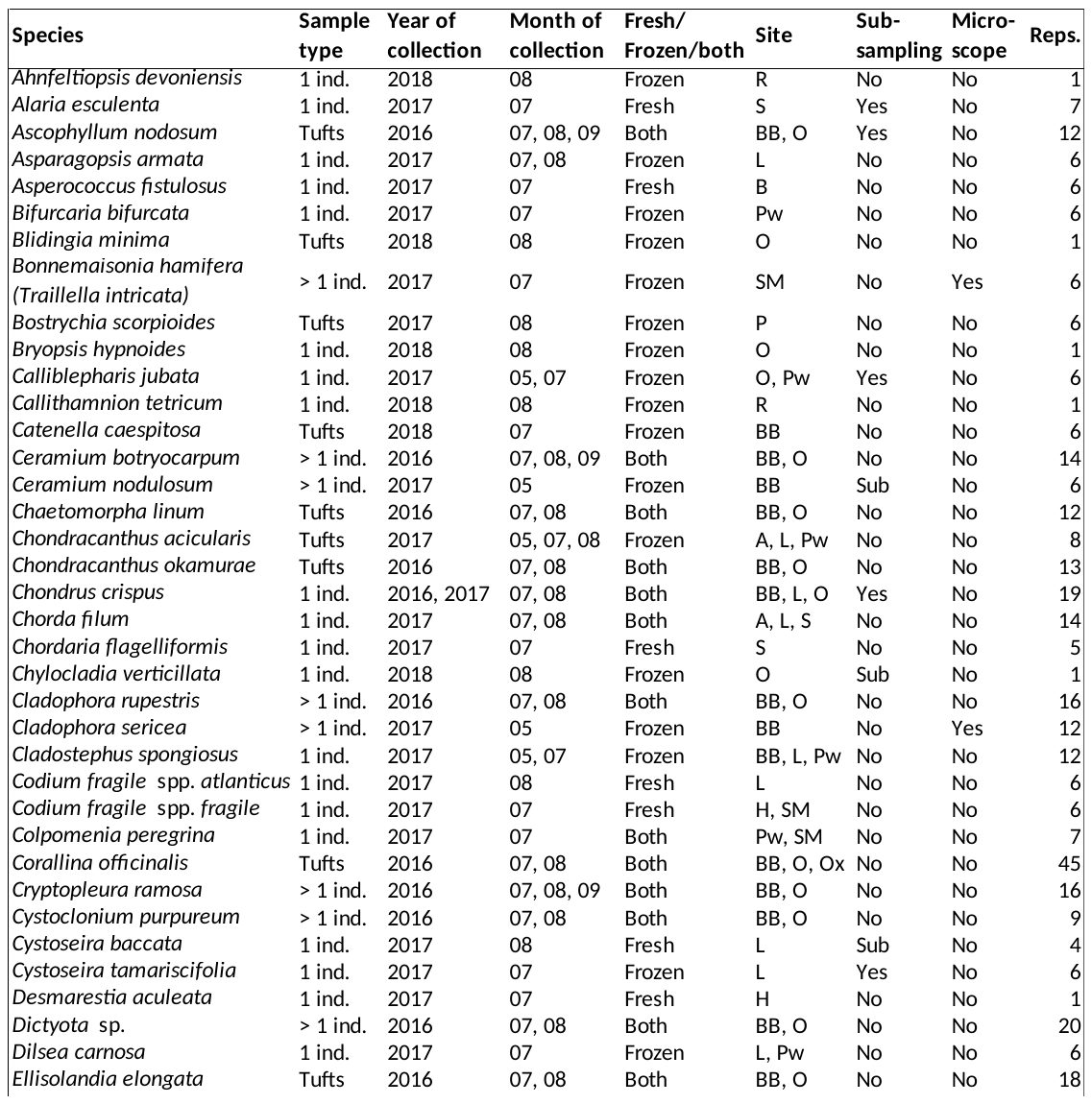

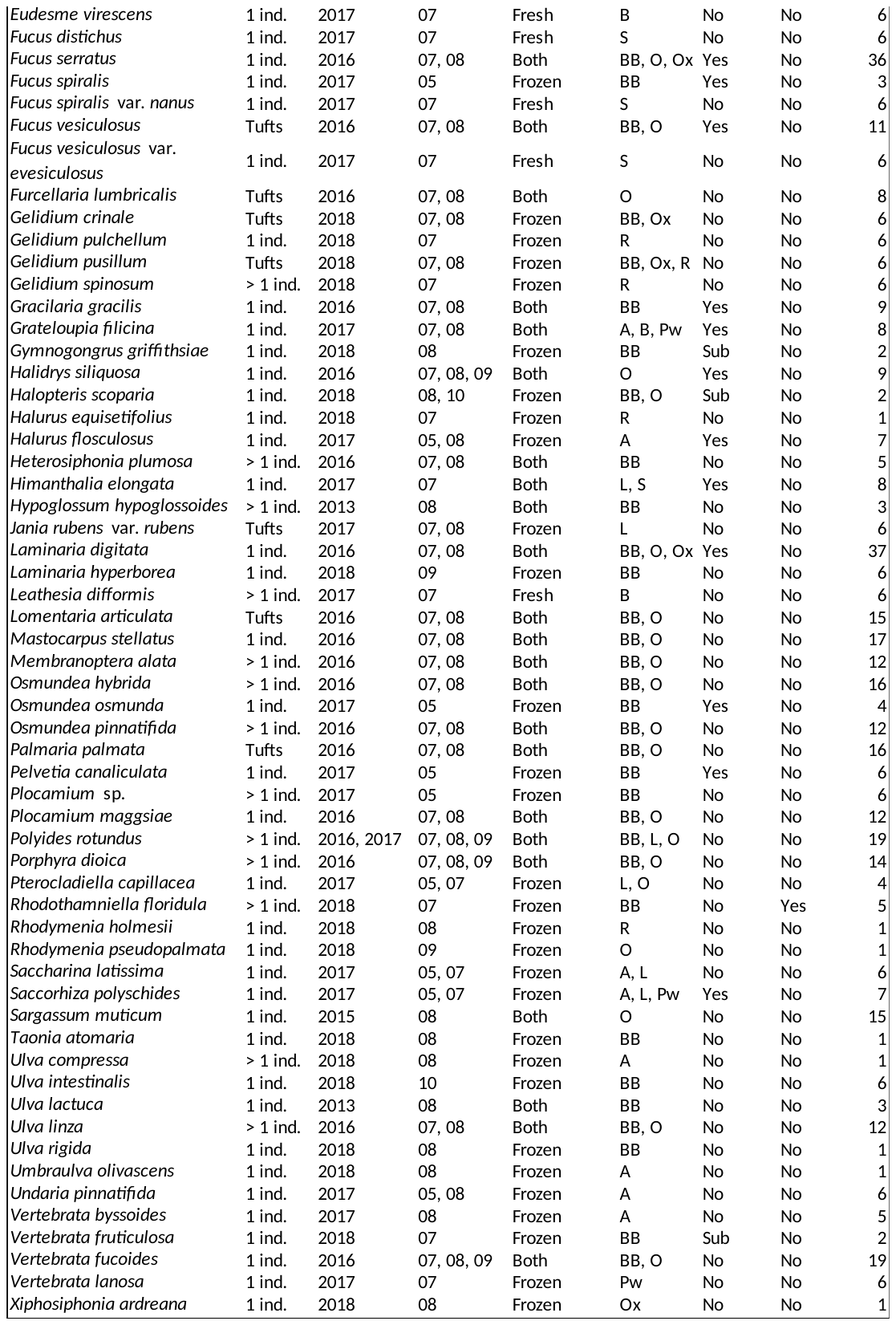

**Table S5. Trait transformations and weightings used in the Gower matrix.** We transformed species trait means to make them follow a distribution that was as close to normality as possible and reduce differences in scale across traits. The traits measured were Specific Thallus Area (STA, mm² g^-1^), the ratio between Surface Area and Volume (SA:V, mm² ml^-1^), thickness (mm), Nitrogen content (N), Thallus Dry Matter Content (TDMC), the ratio between Carbon and Nitrogen content (C:N), Carbon content (C), maximum length, pneumatocysts (YES/NO), aspect ratio, branching order, and the ratio between Surface Area and Perimeter (SA:P). Because some of the traits were correlated, we weighted the Gower matrix on which the Principal Coordinate Analysis is run based on their degree of hypothesised functional redundancy.

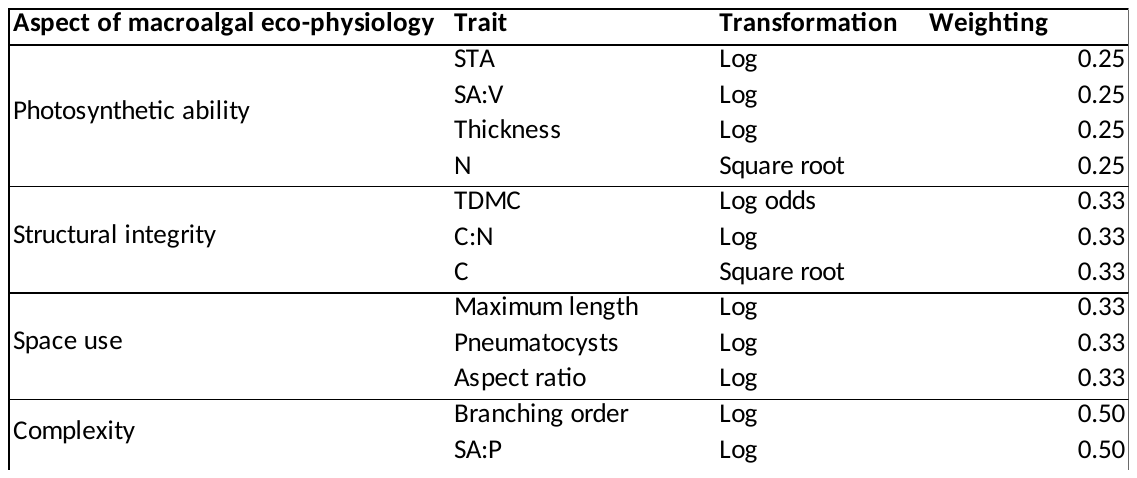

**Table S6. Significant trait correlations and their linearity.** We provide Spearman's *ρ* and level of significance (‘*’ for *P* < 0.05, ‘**’ for *P* < 0.01, and ‘***’ for *P* < 0.001) for all significant correlations between transformed species-level traits (Table S5). The traits measured are Specific Thallus Area (STA, mm² g^-1^), the ratio between Surface Area and Volume (SA:V, mm² ml^-1^), thickness (mm), Nitrogen content (N), Thallus Dry Matter Content (TDMC), the ratio between Carbon and Nitrogen content (C:N), Carbon content (C), maximum length, pneumatocysts (YES/NO), aspect ratio, branching order, and the ratio between Surface Area and Perimeter (SA:P; Table S5). We also visually assessed the linearity of the correlations from pair plots. Many of the traits were rather strongly correlated and some correlations were clearly not linear.

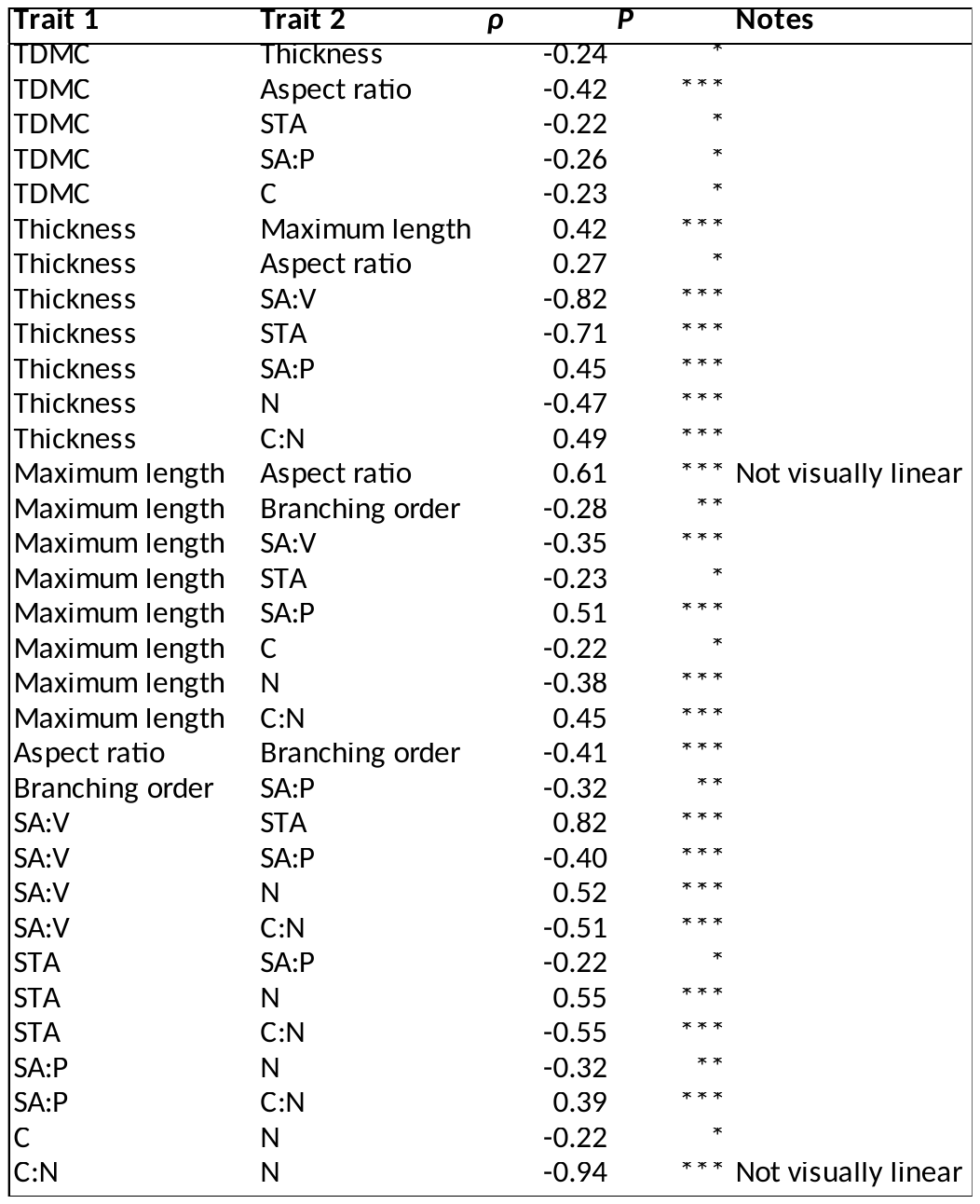

**Table S7. Differences in species composition across grouping methods.** The proportion (%) of species that were allocated inconsistently across grouping methods is given for each of Littler and Littler’s groups and overall. We compared Littler and Littler’s groups to their analogues in Steneck and Dethier’s classification; the two *post hoc* clustering approaches undertaken (agglomerative Hierarchical Agglomerative Clustering or HCA and *k*-medoids) using five and seven clusters; Steneck and Dethier’s five- and seven-group classification; and two common classifications of vertical space use. For example, ca. 8% of species belonging to the ‘sheet’ group of Littler and Littler’s scheme and the analogous ‘foliose’ group of Steneck and Dethier’s scheme were classified inconsistently across the two emergent classifications.

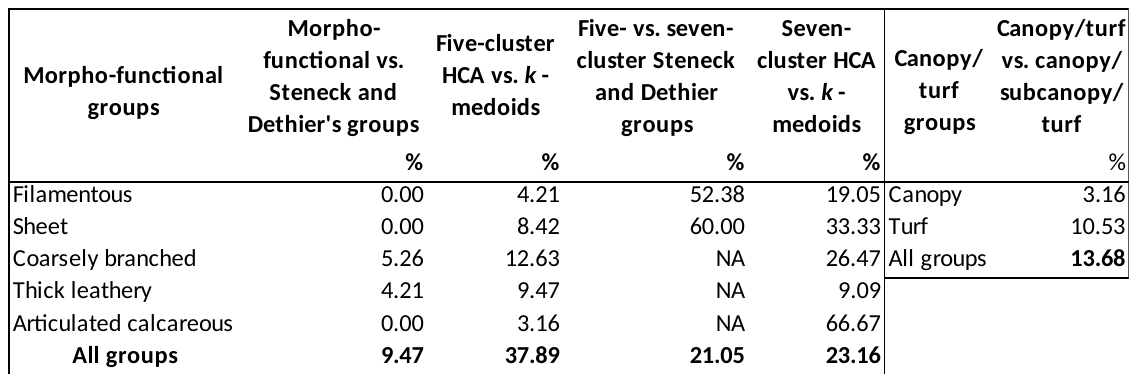

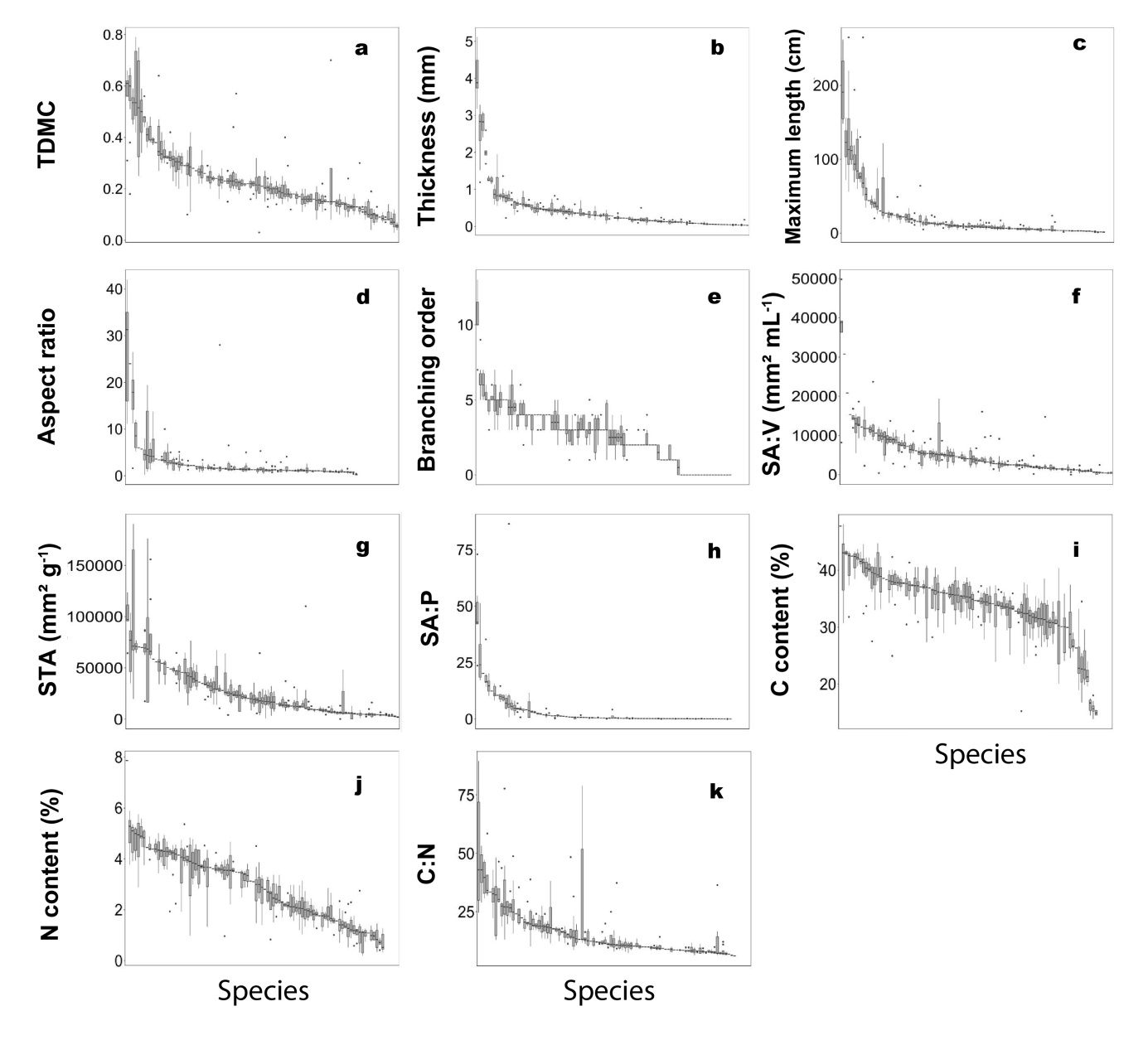

**Figure S1. Intra- and interspecific variability among the continuous functional traits measured.** Distribution of trait attributes are displayed across the 95 macroalgal species screened and for the eleven continuous functional traits measured. *Chorda filum* was excluded from plot d. as its aspect ratio values were much higher than the rest of the species (ranging from 293 to 885, median = 545, S.D. = 205). A series of Kruskal-Wallis tests performed on each functional trait were all highly significant (TDMC: *χ*² = 434.77, d.f. = 94, *P* < 0.001; thickness: *χ*² = 487.62, d.f. = 94, *P* < 0.001; maximum length: *χ*² = 552.63, d.f. = 91, *P* < 0.001; aspect ratio: *χ*² = 276.47, d.f. = 80, *P* < 0.001; branching order: *χ*² = 424.58, d.f. = 88, *P* < 0.001; SA:V: *χ*² = 447.85, d.f. = 94, *P* < 0.001; STA: *χ*² = 441.76, d.f. = 94, *P* < 0.001; SA:P: *χ*² = 467.45, d.f. = 88, *P* < 0.001; C: *χ*² = 331.43, d.f. = 89, *P* < 0.001; N: *χ*² = 399.02, d.f. = 89, *P* < 0.001; C:N: *χ*² = 394.35, d.f. = 89, *P* < 0.001).

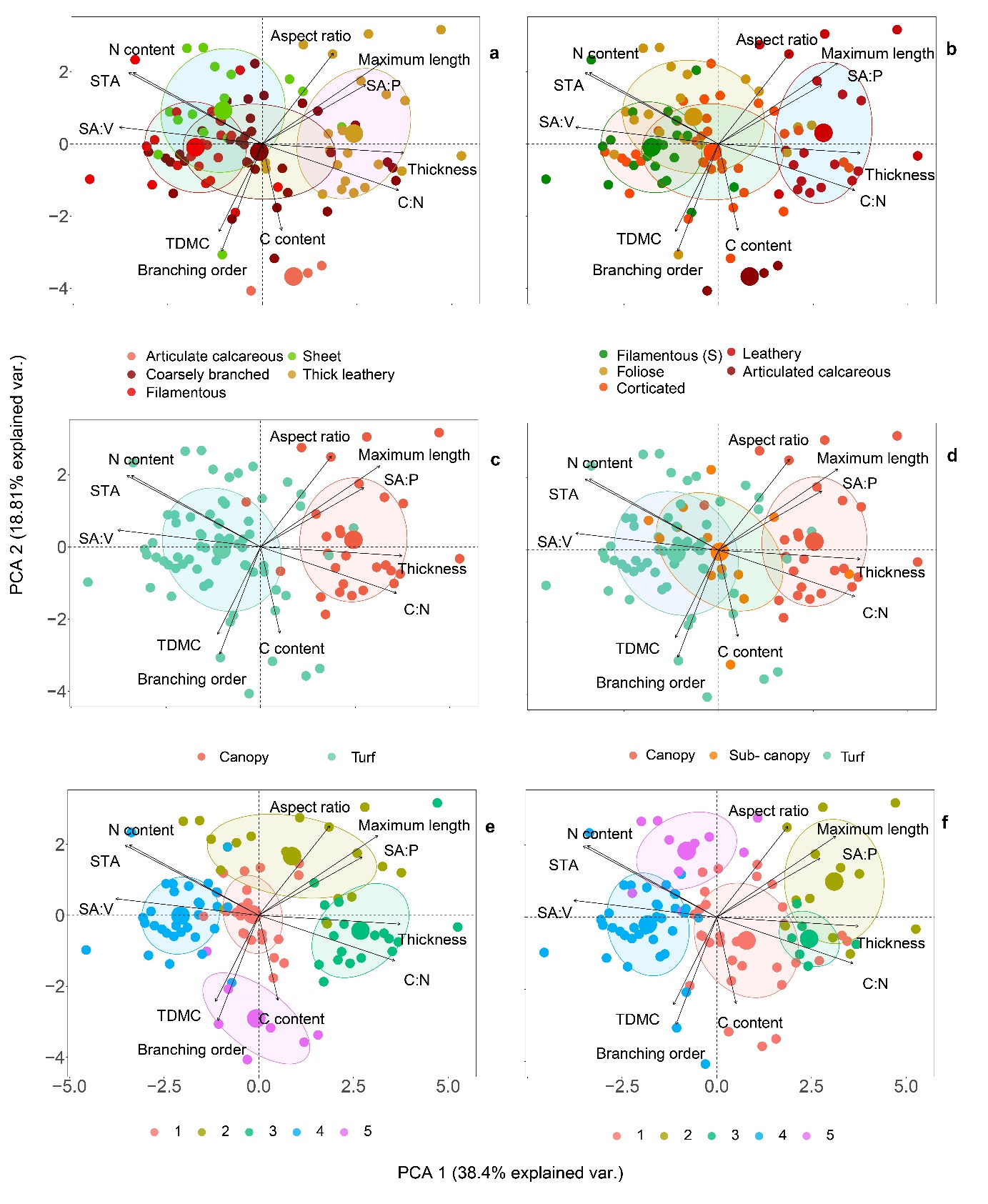

**Figure S2. Distribution of functional groups along the first two principal component axes.** Group distribution in trait space as represented by the first two axes of a principal component analysis (PCA 1 and PCA 2) is given for: (a) Littler and Littler’s ‘functional-form’ model, (b) Steneck and Dethier’s classification, (c) canopy vs. turf, (d) canopy/subcanopy/turf, and emergent groups yielded by *post hoc* clustering of the species using (e) agglomerative Hierarchical Clustering Analysis (HCA) and (f) the divisive *k*-medoids method. Concentration ellipses (50%) are represented.

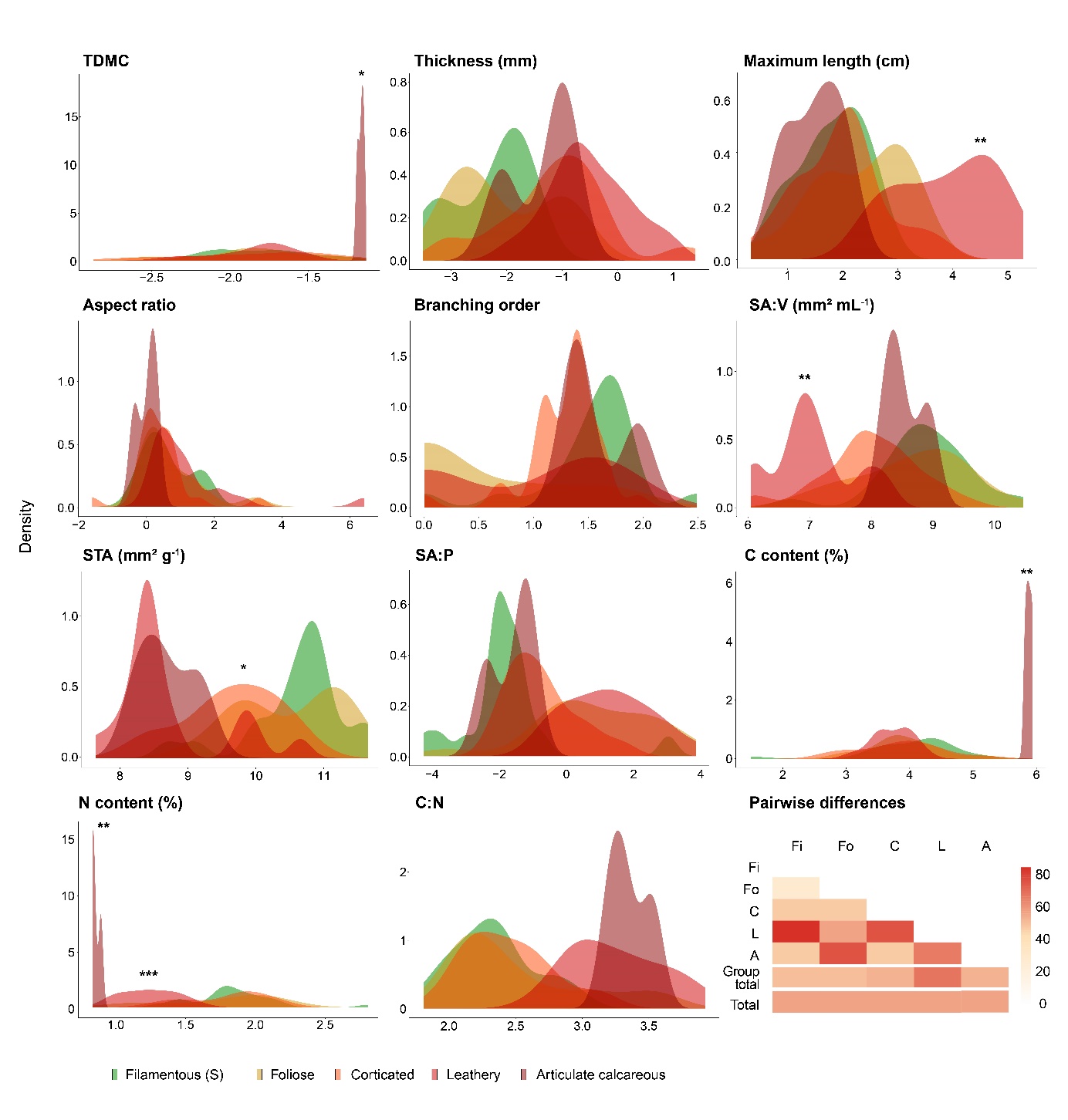

**Figure S3. Distribution of species-level traits among Steneck and Dethier’s groups.** Values of the 11 continuous functional traits are transformed species-level means across the 95 macroalgal species screened (Table S5). Groups that are significantly different from all others are marked by asterisks (pairwise Wilcoxon rank sum test; ‘*’: *P* < 0.05, ‘**’: *P* < 0.01, ‘***’: *P* < 0.001; lowest common *P* is shown). The proportion of significant differences among pairwise comparisons is shown as a heatmap for each group (group total) and overall (total; ‘Fi’ stands for filamentous (S), ‘Fo’ for foliose, ‘C’ for corticated, ‘L’ for leathery, and ‘A’ for articulated calcareous); see Fig. S5 for pairwise differences among groups for each individual trait.

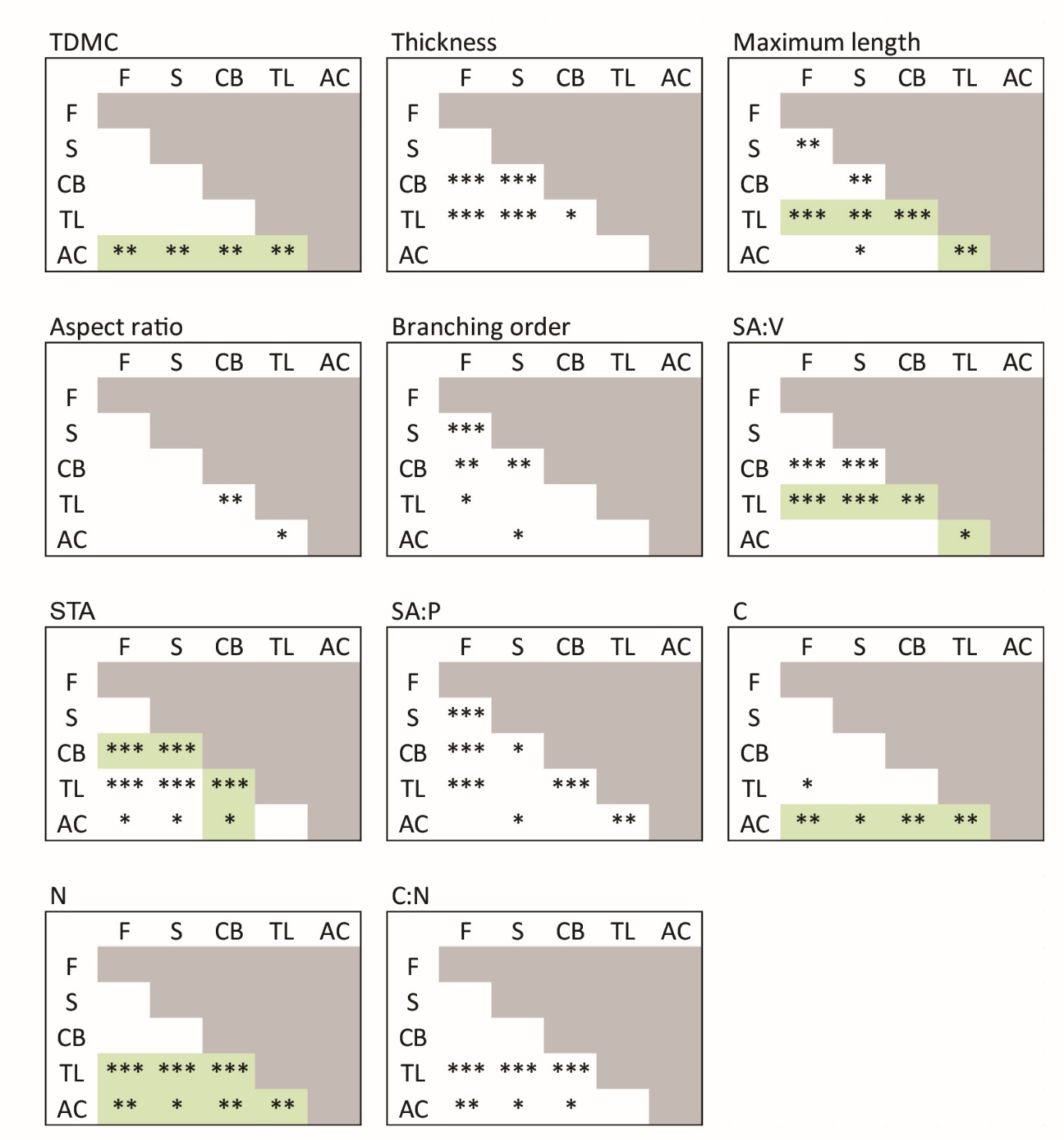

**Figure S4. Pairwise differences among Littler and Littler’s groups.** Significant differences are indicated by asterisks (pairwise Wilcoxon rank sum test; ‘*’: *P* < 0.05, ‘**’: *P* < 0.01, ‘***’: *P* < 0.001). Groups that are significantly different from all others are highlighted in green. Group abbreviations are as follows: ‘AC’ for articulated calcareous, ‘CB’ for coarsely branched, ‘F’ for filamentous, ‘S’ for sheet, and ‘TL’ for thick leathery.

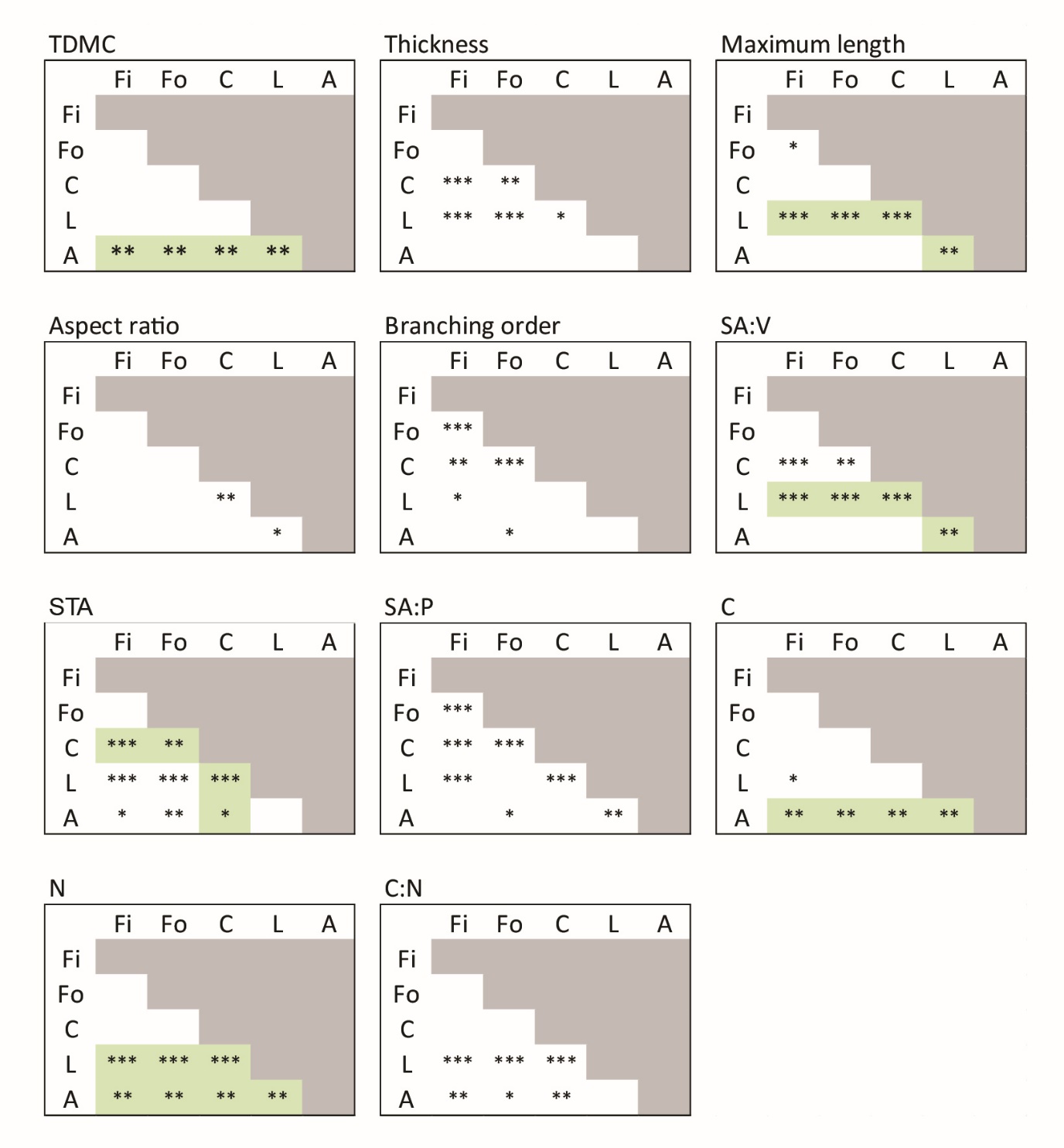

**Figure S5. Pairwise differences among Steneck and Dethier’s groups.** Significant differences are indicated by asterisks (pairwise Wilcoxon rank sum test; ‘*’: *P* < 0.05, ‘**’: *P* < 0.01, ‘***’: *P* < 0.001). Groups that are significantly different from all others are highlighted in green. Group abbreviations are as follows: ‘A’ for articulate calcareous, ‘C’ for corticated, ‘Fi’ for filamentous, ‘Fo’ for foliose, and ‘L’ for leathery.

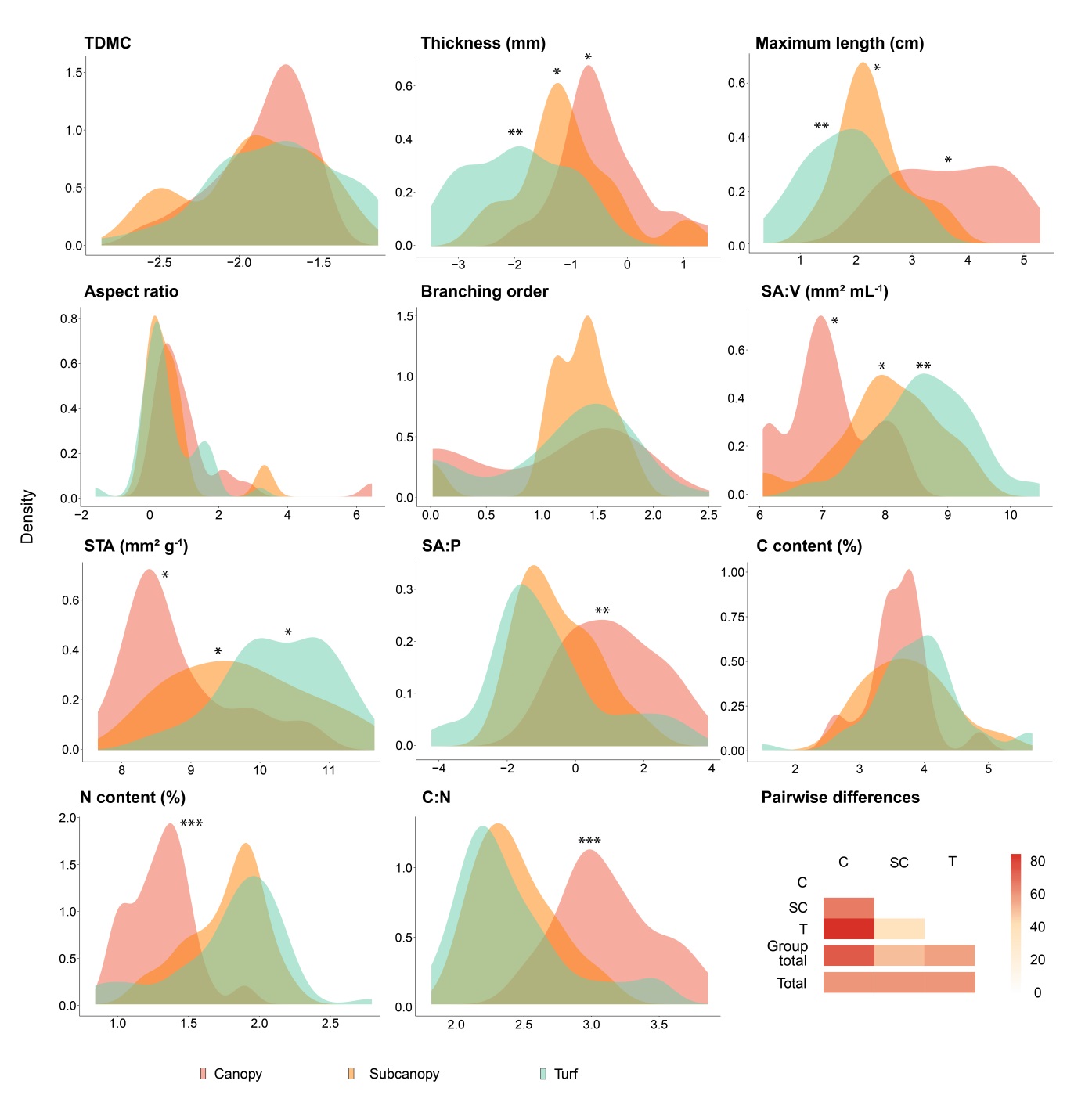

**Figure S6. Distribution of species-level traits among a common three-level classification of vertical space use.** Values of the 11 continuous functional traits are transformed species-level means across the 95 macroalgal species screened (Table S5). Groups that are significantly different from all others are marked by asterisks (pairwise Wilcoxon rank sum test; ‘*’: *P* < 0.05, ‘**’: *P* < 0.01, ‘***’: *P* < 0.001; lowest common *P* is shown). The proportion of significant differences among pairwise comparisons is shown as a heatmap for each group (group total) and overall (total; ‘C’ stands for canopy, ‘SC’ for subcanopy, and ‘T’ for turf); see Fig. S7 for pairwise differences among groups for each individual trait.

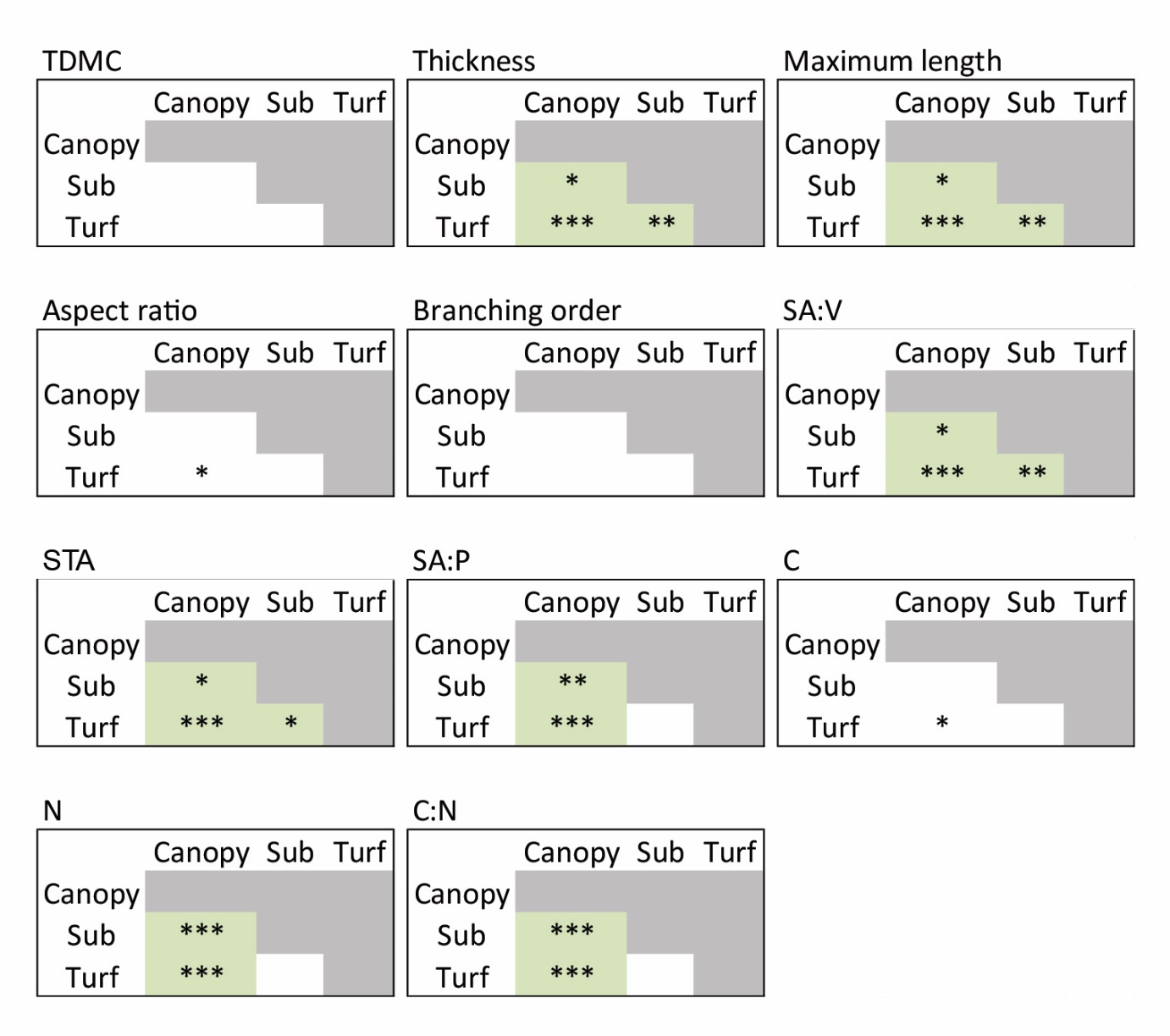

**Figure S7. Pairwise differences among the groups of a common three-level classification of macroalgal vertical space use.** Significant differences are indicated by asterisks (pairwise Wilcoxon rank sum test; ‘*’: *P* < 0.05, ‘**’: *P* < 0.01, ‘***’: *P* < 0.001). Groups that are significantly different from all others are highlighted in green. The group ‘subcanopy’ is abbreviated as ‘sub’.

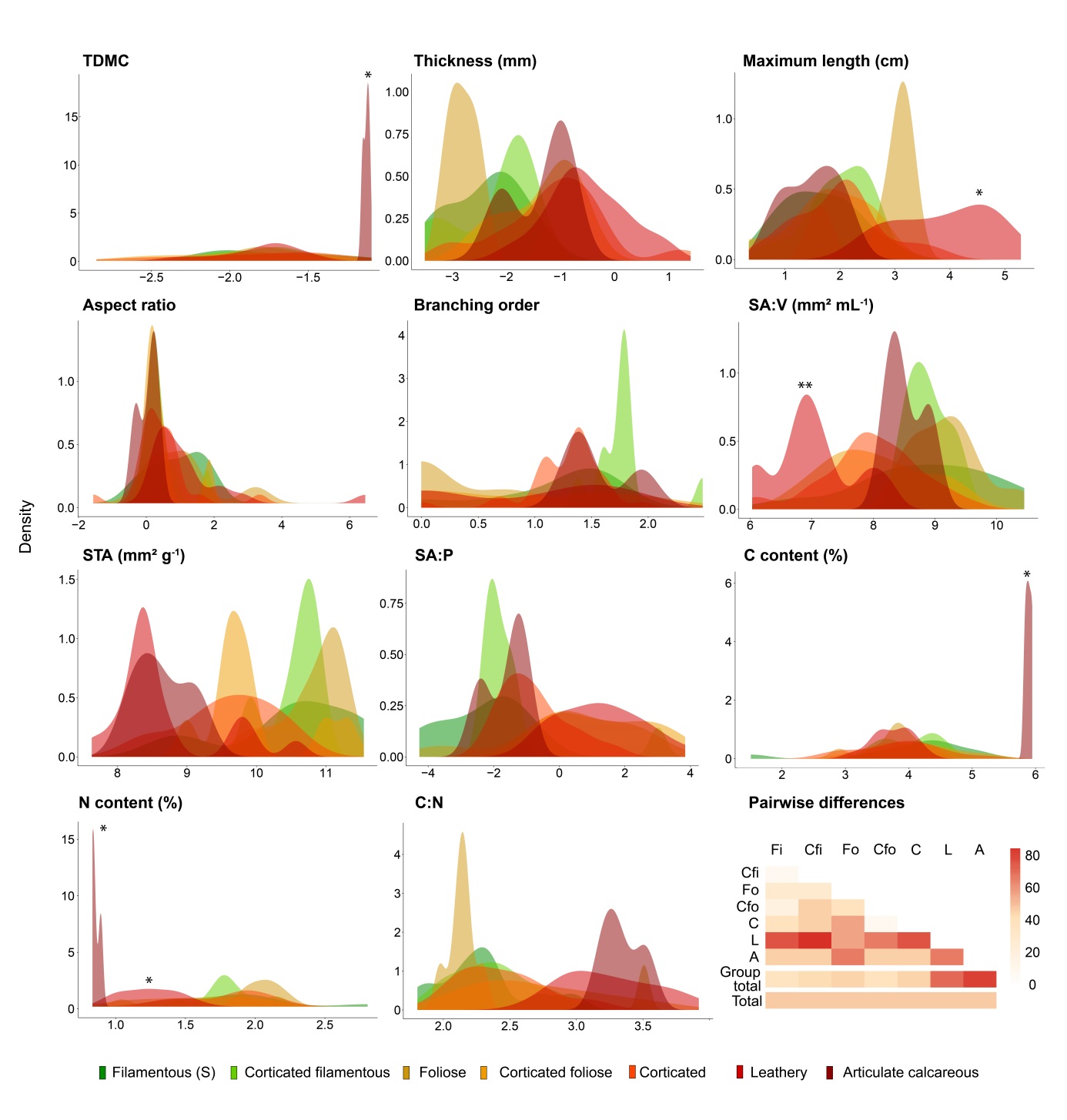

**Figure S8. Distribution of species-level traits among the groups established by Steneck and Dethier’s seven-cluster classification.** Values of the 11 continuous functional traits are transformed species-level means across the 95 macroalgal species screened (Table S5) and are displayed as smoothed density curves. Groups that are significantly different from all others are marked by asterisks (pairwise Wilcoxon rank sum test; ‘*’: *P* < 0.05, ‘**’: *P* < 0.01; lowest common *P* is shown The proportion of significant differences among pairwise comparisons is shown as a heatmap for each group (group total) and overall (total; ‘Fi’ stands for Filamentous, ‘Cfi’ for Corticated filamentous, ‘Fo’ for Foliose, ‘Cfo’ for Corticated Foliose, ‘Co’ for Corticated, ‘L’ for Leathery and ‘A’ for Articulate calcareous); see Fig. S9 for pairwise differences among groups for each individual trait.

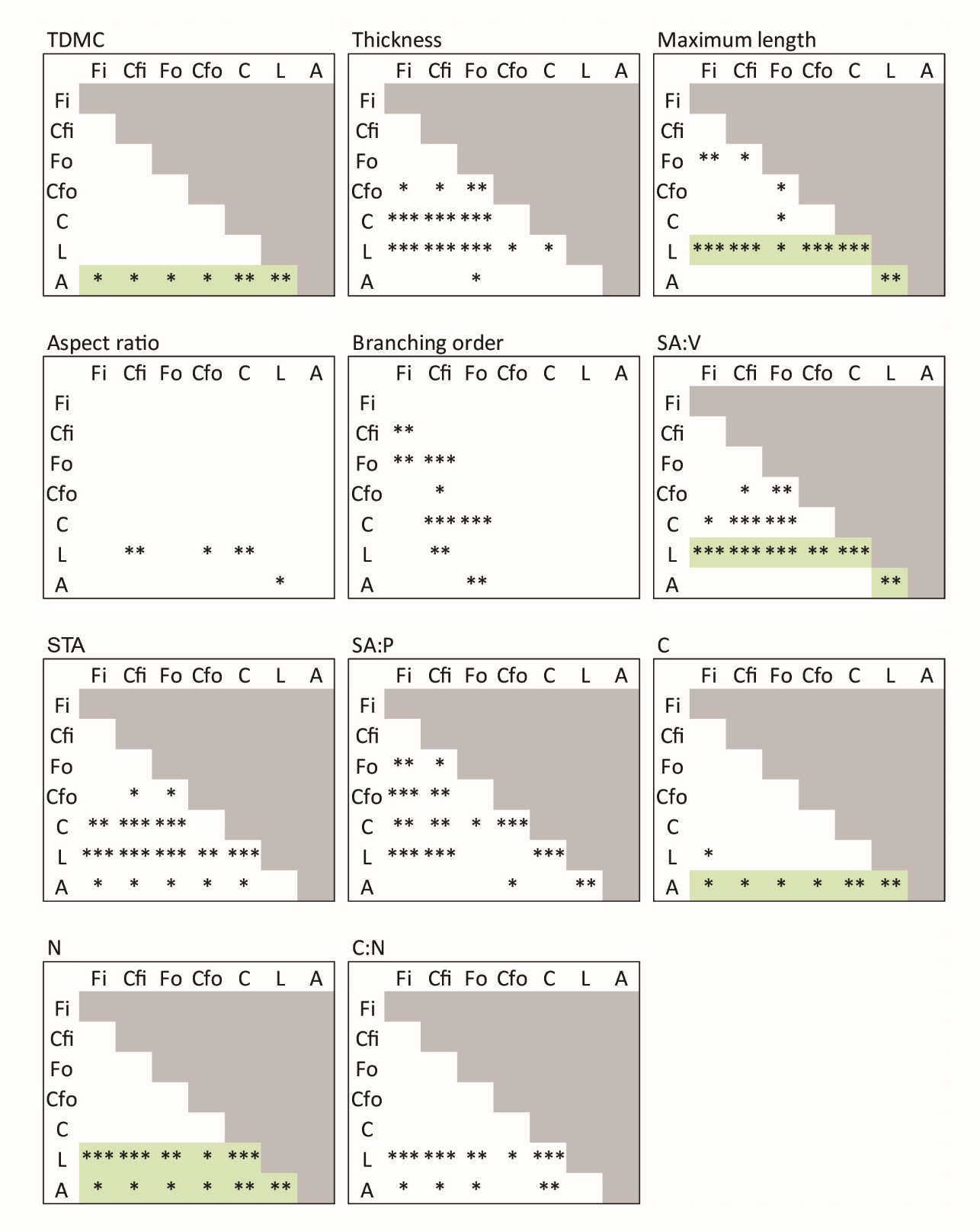

**Figure S9. Pairwise differences among the groups of Steneck and Dethier’s seven-cluster classification.** Significant differences are indicated by asterisks (pairwise Wilcoxon rank sum test; ‘*’: *P* < 0.05, ‘**’: *P* < 0.01, ‘***’: *P* < 0.001). Groups that are significantly different from all others are highlighted in green. Group abbreviations are as follows: ‘Fi’ for Filamentous, ‘Cfi’ for Corticated filamentous, ‘Fo’ for Foliose, ‘Cfo’ for Corticated Foliose, ‘Co’ for Corticated, ‘L’ for Leathery and ‘A’ for Articulate calcareous.

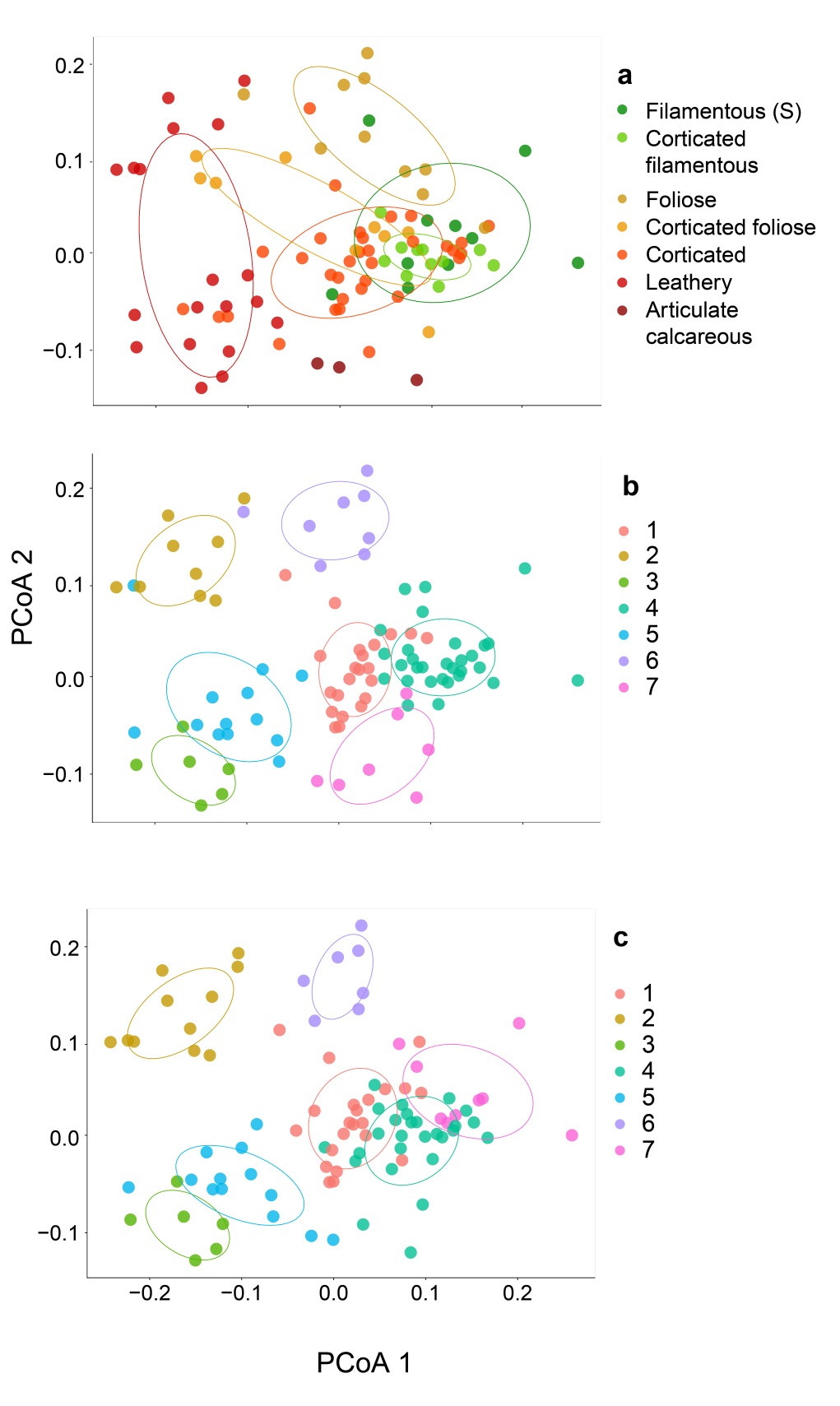

**Figure S10. Distribution of seven-cluster functional groups along the first two principal coordinate axes.** Group distribution in trait space as yielded by a Principal Coordinate Analysis (PCoA) is given for: (a, b) Steneck and Dethier’s detailed classification and emergent groups yielded by *post hoc* clustering of the species using (c, d) agglomerative Hierarchical Clustering Analysis (HCA) and (e, f) the divisive *k*-medoids method. Confidence ellipses (50%) are represented, assuming a multivariate normal distribution.

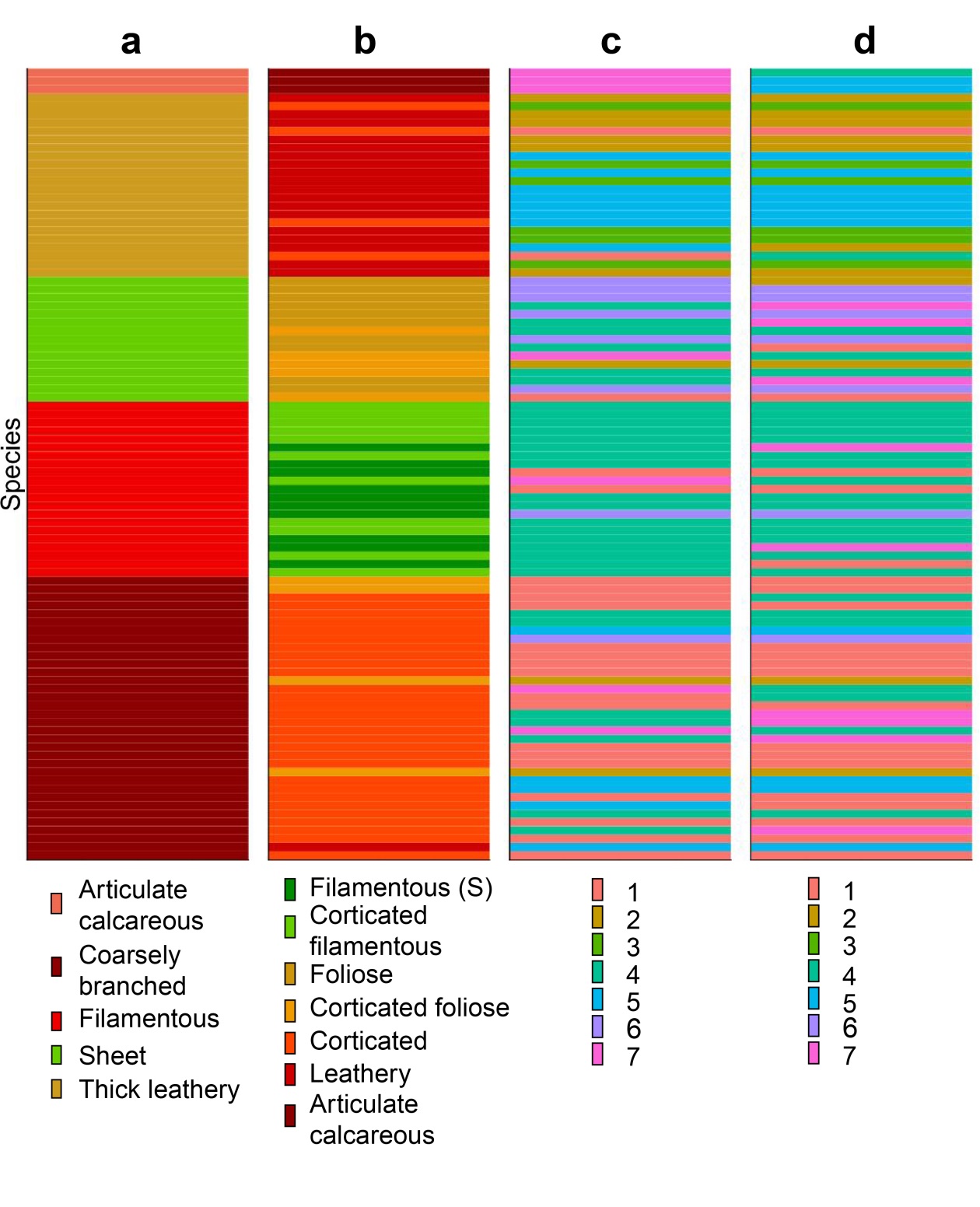

**Figure S11. Species re-allocation across seven-cluster grouping approaches.** Species composition is given for (a) Littler and Littler’s ‘functional-form’ model (ordered alphabetically), (b) Steneck and Dethier’s classification (ordered by degree of cortication), and emergent groups created by *post hoc* clustering of our data using (c) agglomerative Hierarchical Clustering Analysis (HCA) and (d) the divisive *k*-medoids method. Within each grouping approach, stacked horizontal bars correspond to individual species. The order of individual species remains consistent across columns and is initially ordered according to Littler and Littler’s groups. Tracking individual species from left to right shows re-classification across grouping methods.
